# G1/S transcription factors assemble in discrete clusters that increase in number as cells grow

**DOI:** 10.1101/574772

**Authors:** Labe Black, Sylvain Tollis, Guo Fu, Jean-Bernard Fiche, Savanna Dorsey, Jing Cheng, Ghada Ghazal, Stephen Notley, Benjamin Crevier, Jeremy Bigness, Marcelo Nollmann, Mike Tyers, Catherine A. Royer

**Affiliations:** Department of Biological Sciences, Rensselaer Polytechnic Institute, Troy NY 12180 USA; Institute for Research in Immunology and Cancer, Université de Montréal, Montréal, Quebec, H3T 1J4 Canada.; Wellcome Trust Centre for Cell Biology, Institute of Cell Biology, School of Biological Sciences, The University of Edinburgh, Edinburgh EH9 3JR, United Kingdom; Centre de Biochimie Structurale, CNRS UMR5048, INSERM U1054, Université de Montpellier, 29 Rue de Navacelles, 34090 Montpellier, France

**Author notes:** These authors contributed equally to this work (Labe Black, Sylvain Tollis, Mike Tyers, Catherine A. Royer).

## Abstract

The spatio-temporal organization of transcription factor (TF)-promoter interactions is critical for the coordination of transcriptional programs. In budding yeast, the main G1/S transcription factors, SBF and MBF, are limiting with respect to target promoters in small G1 phase cells and accumulate as cells grow, raising the question of how SBF/MBF are dynamically distributed across the G1/S regulon. Super-resolution Photo-Activatable Localization Microscopy (PALM) mapping of the static positions of SBF/MBF subunits revealed that 85% were organized into discrete clusters containing ∼8 copies regardless of cell size, while the number of clusters increased with growth. Stochastic simulations with a mathematical model based on co-localization of promoters in clusters recapitulated observed cluster behavior. A prediction of the model that SBF/MBF should exhibit both fast and slow dynamics was confirmed in PALM experiments on live cells. This spatio-temporal organization of the TFs that activate the G1/S regulon may help coordinate commitment to division.

## Introduction

Commitment to cell division occurs in late G1 phase, an event termed Start in budding yeast (Hartwell et al., 1974; Johnston et al., 1977). Start depends on an extensive G1/S transcriptional regulon comprised of ∼200 genes that function in macromolecular biosynthesis, bud emergence, DNA replication, SPB duplication and other critical processes. The G1/S transcriptional program is controlled by two master transcription factor (TF) complexes, SBF and MBF, made up of one DNA binding subunit, Swi4 and Mbp1, respectively, and a common activator subunit, Swi6 (Koch et al., 1993). SBF and MBF recognize specific sites in G1/S promoter regions, called SCB and MCB sites, with some degree of overlapping specificity (Bean et al., 2005; Iyer et al., 2001; Koch et al., 1993). ChipSeq experiments have delineated Swi4, Mbp1 and Swi6 binding sites in the genome (Iyer et al., 2001; Lee et al., 2002; Park et al., 2013; Simon et al., 2001), although the agreement between these various studies is only partial (Ferrezuelo et al., 2010).

Based on recent Swi6 ChipSeq data, bioinformatics approaches have been used to map the Swi6 target sites onto a 3D model of the budding yeast G1 phase genome (Capurso et al., 2016; Duan et al., 2010; Park et al., 2013). This model predicted functional 3D hotspots for Swi6 binding, in particular the *MSB2* and *ERG11* genes. A combination of ChipSeq and chromatin capture data suggests many transcription factors in budding yeast, including Swi4 and Swi6, have targets sites that cluster in space (Ben-Elazar et al., 2013; Duan et al., 2010; Eser et al., 2017). Swi4 and Swi6 have been shown to be associated with highly transcriptionally active gene clusters (Tsochatzidou et al., 2017). While these domains seem to separate regions of similarly timed replication origins, their relation to the timing of the Start transition has not been characterized. Despite the strong inference of TF clustering from these studies, the spatial and temporal organization of the G1/S TFs and their target sites has not been directly observed.

Here, we have used a super-resolution method, Photo-Activatable Localization Microscopy (PALM) (Betzig et al., 2006; Rust et al., 2006) to map the static and dynamic positions of fusions of Swi4, Mbp1 and Swi6 with the photoactivatable protein mEos3.2 (Zhang et al., 2012) expressed from their natural loci in fixed and live budding yeast cells. The resultant PALM images of fixed cells provided 2D projections of the 3D organization of these proteins in the nucleus. We found that the TFs organize into clusters of ∼8 monomers (4 dimers) that range in number from ∼5 in small cells to ∼30 in large cells. Given that, throughout most of G1, SBF/MBF copy numbers are limiting with respect to the ∼200 G1/S promoters (Dorsey et al., 2018), the observed SBF/MBF clustering strongly suggests close spatial proximity of several promoter sites within each cluster. While the number of clusters increased with cell size, the number of molecules per cluster was independent of cell size. This increase in TF cluster number was in overall in agreement with our previous observations of an increase in TF copy number as cells grow (Dorsey et al., 2018). A mathematical model and Monte Carlo computer simulations of TF clustering constrained by these observations and simple biophysical assumptions predicted that TFs should alternate between a highly confined state, in which they are trapped within G1/S promoter clusters, and a highly dynamic state, in which they hop rapidly between clusters. Live cell single particle tracking (spt)-PALM verified the prediction of distinct slow sub-diffusive and fast diffusive dynamic modes for these factors. Overall, these results suggest that the promoters of the G1/S regulon are spatially organized into discrete clusters that are successively titrated by increasing TF copy number as cells grow.

## Results

### The G1/S transcription factors are clustered in yeast nuclei

Super-resolution PALM images of mEos3.2 fusions of Swi4, Mbp1 and Swi6 in single nuclei from fixed cells grown on rich (SC+2% glucose) medium revealed non-homogeneous distributions for each protein (Figure 1, Figures 1-Supplemental Figures 1 and 2). No size phenotype was observed for these strains, as with our prior studies on strains expressing GFP fusions of these factors (Dorsey et al., 2018), indicating that these crucial TFs retain their function when fused to the fluorescent proteins. The super-resolution detection images (Figures 1A-D, Figures 1-Supplemental Figures 1A-D and 2 A-D) (resolution ∼25 nm) result from the super-position of all detections of all molecules over the ∼30,000-40,000 frames (30 ms exposure time/frame) acquired for a given field of view (FOV). However, since each protein was cross-linked by fixation and thus immobile, and was detected multiple times in successive frames (up to 100), it is possible to average its mean position over multiple detections to obtain a molecular (Betzig et al., 2006) as opposed to detection image, in which the average position of each individual TF molecule is represented (Figure 1 E-G; Figures 1-Supplemental Figures 1E-G and 2 E-G). It is apparent from both the detection and the molecular images that in all cells (small, medium and large) most nuclear Swi6, Mbp1 and Swi4 molecules are organized in discrete clusters of a few molecules. Clusters of the TFs were also observed in cells grown on SC+2% glycerol, a poor carbon source (Figures 1-Supplemental Figure 3). The super-resolution images correspond to 2D representations of 3D objects, since the microscope depth of field (∼500nm) is larger than macromolecular structures. Hence, some degree of clustering could in principle arise from super-position of molecules in different z-planes. Nonetheless, the extensive degree of clustering observed exceeds what may be expected from 2D super-position of randomly distributed molecules in 3D (see simulations below).

**Figure 1.**
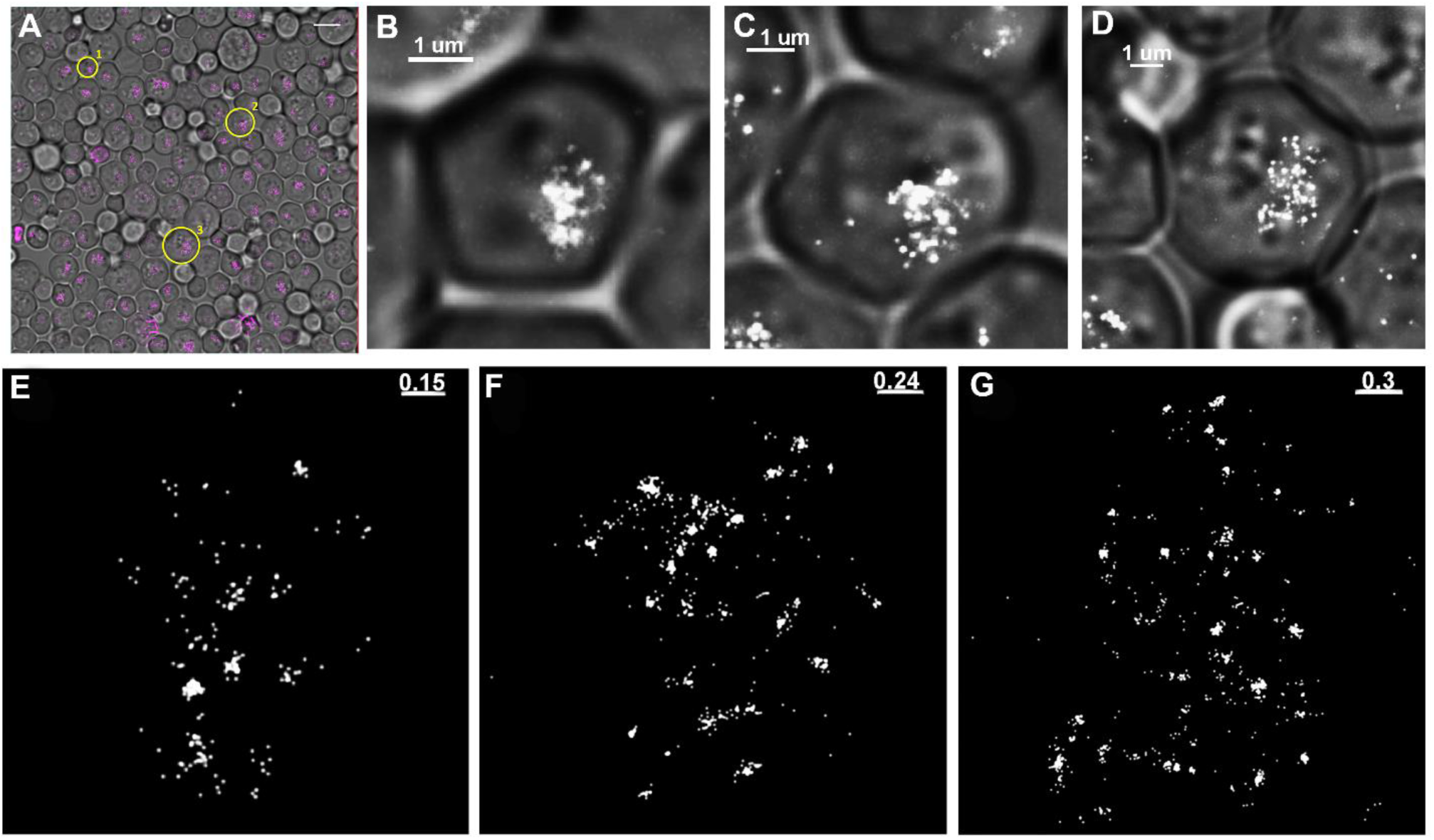
Super-resolution PALM imaging reveals clustering of Swi4-mEos3.2 in fixed budding yeast cells. A) Composite image of the phase contrast and PALM images of a FOV of Swi4-mEos3.2 cells grown in SC+2%glucose. Detection image outputs (pink dots; one for each detection) were obtained with the Thunderstorm plugin from ImageJ. Detection images were not filtered for blinking. High intensity purple in the image corresponds to out of focus beads. Zoomed cells 1-3 are indicated by yellow circles and numbers. Scale bar is 10 μm. B-D) Zoomed composite images of small (1), medium (2) and large (3) cells. Scale bars are 1 μm. E-F) Molecular images of the nuclei of cells 1-3, respectively. Scale bars are 0.15 μm in panel E, 0.24 μm in panel F and 0.30 μm in panel G.

### The number of clusters of G1/S transcription factors increases with cell size while copy number per cluster remains constant

Nuclei were masked, and the number of molecules detected in each nucleus was obtained from analysis of the blinking-corrected molecular images (i.e., Figures 1E-G, Figure 1-Supplemental Figures 1 and 2E-G) as described in the methods section and SI. Given their nuclear size, most cells to the left of the black dashed lines in Figure 2 are expected to be in G1 phase (see Methods, Figure 2-Supplemental Figure 1). The nuclear copy numbers of Swi4, Mbp1 and Swi6 increased with cell size (Figure 2A-C, respectively), consistent with our previously reported size-dependent increase in G1/S copy number determined by Number and Brightness (N&B) fluctuation microscopy (Dorsey et al., 2018). The average number of proteins per nucleus was in reasonably good agreement with values determined by N&B of 50-100 copies in small G1 phase cells and 100-200 in large G1 phase cells (Figure 2-Supplemental Figure 2), although we detected somewhat fewer molecules in the PALM experiments, especially in large cells. This difference is most likely due to the limited depth of field in the PALM experiments (see Methods section). We note that PALM microscopy is not as reliable as N&B for particle counting due to blinking of mEos3.2, imperfect correction thereof (Lee et al., 2012), incomplete activation of mEos3.2 and exclusion of out-of-focus particles.

**Figure 2.**
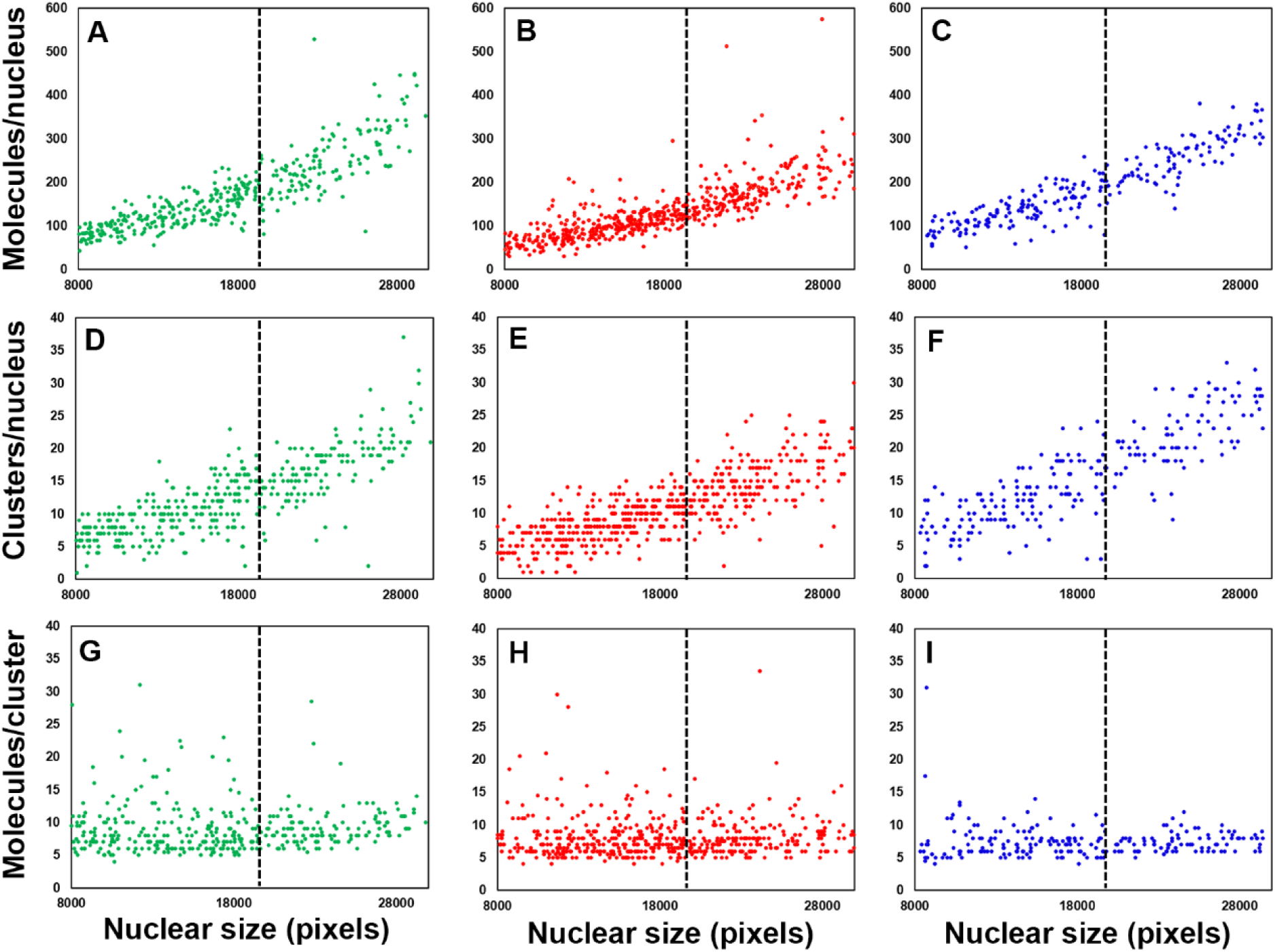
Copy numbers and the number of clusters per cell for Swi4, Mbp1 and Swi6 increase with cell size. A-C) Total number of Swi4-, Mbp1-, and Swi6-mEos3.2 molecules in each nucleus as a function of nuclear size, a proxy for cell size. Each point represents an individual nucleus. D-F) Number of Swi4-, Mbp1-, and Swi6-mEos3.2 clusters per nucleus as a function of cell size. G-I) Average number of Swi4-, Mbp1-and Swi6-mEos3.2 molecules per cluster for each nucleus as a function of cell size. Left-hand panels correspond to Swi4-mEos3.2 (green, 370 cells), middle panels correspond to Mbp1-mEos3.2 (red, 536 cells) and right-hand panels correspond to Swi6-mEos3.2 (blue, 222 cells). Each point represents one nucleus. Nuclear size is given in pixels. To the left of the dashed lines at 20000 pixels the majority of cells are in G1 phase (see Methods and Figure 2 – Supplemental Figure 1).

To quantify the number of clusters in each nucleus and the number of molecules in each cluster, we developed a custom cluster identification algorithm based on a simplification of the OPTICS ranking algorithm (Kriegel et al., 2011). Briefly, our algorithm sorts the list of all detected molecules within each individual nucleus to reorder molecule detections such that, in the final list, each molecule is listed just after (or before) its nearest neighbor in real space (see Methods section). Then it computes the list of distances between each molecule and the next on the list (see Figure 2-Supplemental Figure 3A for an exemplary cell). In order to define and distinguish clusters, a distance threshold (i.e., threshold spike amplitude separating two different clusters) of 10 high-resolution pixels, corresponding to 10 pixels x 3 nm/pix = 30 nm, was chosen (Figure 2-Supplemental Figure 3A, red horizontal line). This threshold value was chosen because the relative change in cluster number with respect to the threshold did not change significantly beyond this critical distance of 30nm (Figure 2-Supplemental Figure 3B). Finally, since Mbp1, Swi4 and Swi6 predominantly occur as dimers (Dorsey et al., 2018), we defined a cluster as a group of molecules larger than at least 2 dimers (i.e., 4 molecules). Unlike previously published cluster detection algorithms (Mazouchi and Milstein, 2015), our algorithm detected the small clusters observed for these proteins, even for the sparse clusters found in small cells. The mEOS3.2 protein has been characterized biophysically and is essentially monomeric (Wang et al., 2014), making it unlikely to be responsible for the clustering of Swi4, Mbp1 and Swi6. As a control we constructed a strain producing from a plasmid, molecules of mEos3.2 fused on both the N-and C-terminus to nuclear localization signal (NLS) peptides to ensure predominantly nuclear localization. As expected, the images of the cells producing the NLS-mEos3.2-NLS construct did not show significant cluster formation (Figure 2 – Supplemental Figure 4). Moreover, unlike the G1/S TFs, cluster analysis of the NLS-mEos3.2-NLS images showed very few wide valleys (corresponding to clusters) in the reordered distance plots (compare Figure 2 – Supplemental Figure 5A to Figure 2 – Supplemental Figure 3A). In addition, the threshold distance dependence of cluster counts exhibited no obvious cutoff value (Figure 2 – Supplemental Figure 5B), compared to the sharp decrease to a constant level observed for the G1/S TFs (Figure 2 – Supplemental Figure 3B), as expected for evenly distributed molecules. These results demonstrated that the clustering observed for the G1/S TFs was not due to any properties (optical or physical-chemical) of the mEos3.2 tag.

For all three TFs, the number of clusters increased with cell size, while the mean number of molecules per cluster was almost invariant with size (Figure 2D-I, Figure 2-Supplemental Figure 6A). Most clusters contained ∼8 molecules (4 dimers), although some were significantly larger (Figure 2G-I; Figure 2-Supplemental Figure 6B). Regardless of cell size, 85% of all molecules were located in clusters, whose lateral extension was in the 30nm-80nm range (Figures 1E-G, Figure 1-Supplemental Figures 1-3E-G and insets). These results suggested that as TF copy number increased with cell growth, TFs form new clusters rather than associating with existing clusters. The number of clusters for each TF in the largest cells reached ∼20-30, much lower than the ∼200 G1/S promoters and the ∼600 target sites across all G1/S promoters (Ferrezuelo et al., 2010; Iyer et al., 2001). Interestingly, Swi6 clusters had about the same average number of molecules as Swi4 or Mbp1 (Figures 2, Figure 2-Supplemental Figure 6), such that the larger number of Swi6 molecules (with respect to Swi4 or Mbp1) was reflected by a larger number of clusters (but significantly smaller than the sum of Swi4 and Mbp1 clusters for any cell size). This observation suggested that most clusters are composed of both MBF and SBF.

Given that Swi4 and Mbp1 copy numbers are only in slight excess with respect to the number of G1/S promoters in large cells at the end of G1-phase (Dorsey et al., 2018), the organization of the TFs into clusters indicates that G1/S promoters might also be clustered, which may help ensure synchronous expression of the G1/S regulon. In this view, most G1/S promoters would be spatially organized into ∼30 clusters of 7-10 promoters each that are successively titrated by G1/S TFs as cells grow. In a contrasting view, the limiting number of Swi4-Mbp1 dimers could in principle partially populate the promoter clusters even in small cells (Dorsey et al., 2018), and newly synthesized molecules would also randomly distribute across all clusters of target sites. This scenario would result in a constant number of clusters that increase in TF copy number per cluster as cells grow. Our data unequivocally support the former model in which most G1/S promoters are spatially organized into clusters that are successively titrated by G1/S TFs as cells grow.

### A quantitative model couples G1/S DNA promoter clusters to TF clusters

We developed a simple mathematical model and used Monte Carlo computer simulations to explore the biophysical parameters that might explain the observed spatial patterns of the G1/S TFs (Figure 3). The goal was to identify the minimum number of parameters with physically reasonable values that could reproduce the experimental clustering observations. The main feature of the model, in addition to the SBF/MBF binding module of our previously published Start model (Dorsey et al., 2018), was the assumption that G1/S promoters form clusters (see below). The SBF/MBF binding module encompasses mass-action kinetic-driven binding of Swi4 and Mbp1 dimers to Swi6 dimers, their binding to DNA, and the converse dissociation reactions. This equilibrium mathematical model was first converted to mass action-like ordinary differential equations and then into stochastic simulations using the Gillespie algorithm, discretized onto a three-dimensional spatial mesh to account for diffusion (David Bernstein, 2005; Jose et al., 2013). Unless otherwise specified, we used a nuclear diffusion coefficient of *D_nuc_* = 2 µm^2^/s (Thattikota et al., 2018). For model equations, assumptions and parameters see Methods and (Dorsey et al., 2018)).

**Figure 3.**
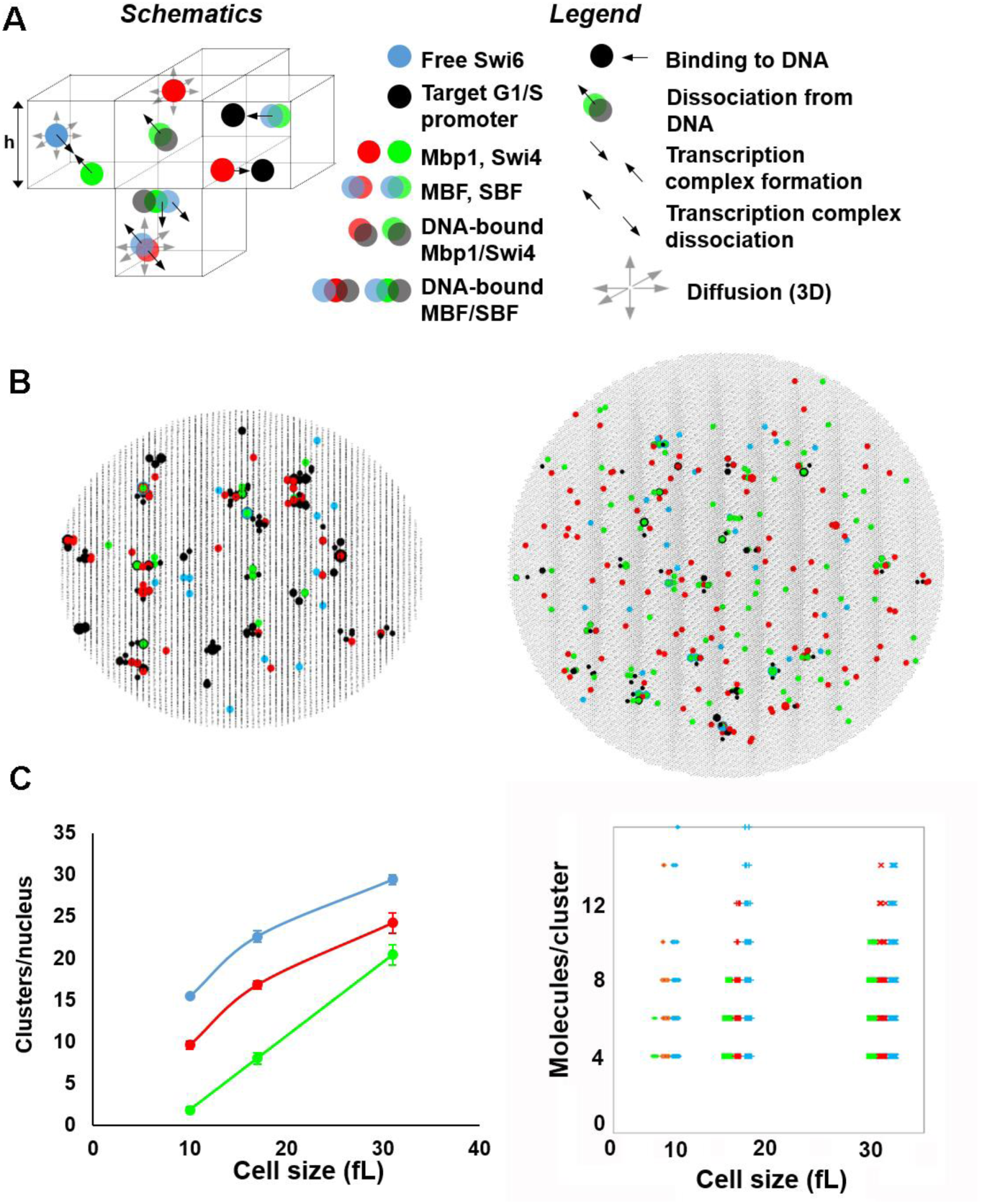
Stochastic modeling predicts Swi4, Mbp1 and Swi6 clustering. A) Schematic of SBF/MBF binding model (left), encompassing mobile Swi4 dimers (green dots), Mbp1 dimers (red dots), Swi6 dimers (blue dots) and immobile G1/S DNA promoters (black dots), moving and interacting in the nucleus discretized in infinitesimal volume elements (cubes separated by thin black lines, only a few of them are shown here). Swi4/Mbp1 can associate with Swi6 to form mobile SBF/MBF, and/or bind (immobile) promoter DNA. For illustrative purposes, the leftmost element shows SBF formation from Swi4 and Swi6 (convergent thick black arrows). The bottommost element shows MBF dissociation into Mbp1 and Swi6 (divergent thick black arrows) and diffusion (thin grey arrows). Also shown is SBF dissociation from DNA. The rightmost element shows Mbp1 and SBF association with 2 promoters within a cluster. All interactions (i.e., promoter DNA binding and dissociation, complex formation and dissociation, diffusion) accounted for in the model are indicated (right). The corresponding propensities (in s^-1^) were derived from (D. Bernstein, 2005) and are listed below for all five reaction types respectively: 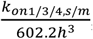, 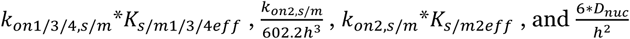, where h is the mesh size, and the reaction “on” rates *k_on_* (and the equilibrium constants *K* such that *k_off_* = *K* ∗ *k_on_*) are defined in Methods and SI.. B) 2-D projection of the 3-dimensional output of a typical simulation showing clusters of Swi4 dimers (green dots), Mbp1 dimers (red), Swi6 dimers (blue) and G1/S DNA promoters (black dots) in small (10fL, left) and large (31.5fL, right) cells. C) (left) Number of Swi4 (green), Mbp1 (red) and Swi6 (cyan) clusters per nucleus (vertical axis) as a function of the size of simulated cells (horizontal axis). Data points indicate the cluster number averaged over ten independent simulations for each cell size, while error bars indicate the standard error on the mean. (Right) Scatter plot showing the number of Swi4 (green), Mbp1 (red) and Swi6 (cyan) molecules per cluster (vertical axis) as a function of the size of simulated cells (horizontal axis). Data points represent individual clusters and were gathered from ten independent simulations for each cell size.

SBF/MBF dimer copy number values as a function of cell size were taken from our previous determination by Number and Brightness microscopy (Dorsey et al., 2018). The concentrations of Mbp1 and Swi6 were found previously to be 110 and 150 nM in G1 cells of all sizes, such that the dimeric copy numbers of 42 and 57, respectively for Mbp1 and Swi6 in small cells increased ∼3-fold to 131 and 178 in large cells. Swi4 concentration was much lower in small cells, 50 nM (dimeric copy number 15) and doubled as cells grow in G1, leading to a dimeric copy number of 109 in large cells (Dorsey et al., 2018). These parameters ensured stable and predominant formation of DNA-bound SBF and MBF complexes, confirming that the equilibrium regime previously predicted (Dorsey et al., 2018) is reached kinetically when molecular noise is accounted for (Figure 3-Supplemental Figure 1A, B). This minimal model is based on the assumption that the ∼200 G1/S promoters are pre-organized and randomly distributed across 35 clusters (yielding typically 4-12 promoters per cluster). It successfully predicts the formation of TF clusters (Figure 3B). Importantly spontaneous TF cluster formation was not observed in simulations when G1/S promoters were not pre-clustered (using identical simulation parameters) (Figure 3-Supplemental Figure 2).

In our simulations and in agreement with our experimental observations, a substantial fraction of promoter clusters was free from binding of any TF in small cells (Figure 3B left and Figure 3 – Supplemental Figure 3A, black dots). In contrast, in larger cells, close to the critical size at the end of G1-phase, most if not all clusters and promoters were bound with either fully formed SBF/MBF or Swi4 or Mbp1 dimers (Figure 3B, right). We computed cluster statistics by counting the number of particles of each type within each promoter cluster (retaining clusters with ≥ 4 molecules, i.e. 2 dimers, to compare with our experiments) across 10 independent simulations for each cell size. The number of Swi4, Mbp1 and Swi6 clusters increased from ∼5-10 in small cells to ∼10-15 for Swi4 and Mbp1 and ∼20 for Swi6 in large cells (Figure 3C, left). The number of molecules per cluster in the model was between 4 and 12 regardless of cell size, with no significant size dependence (Figure 3C, right), in reasonable agreement with our experimental observations (Figures 2, Figure 2-Supplemental Figure 6). Of note, this distribution mirrored the multinomial distribution of promoter cluster size that arose from the random distribution of G1/S promoters across clusters.

We then asked whether clustering could influence TF residence times on each promoter. Ranking all G1/S promoters in simulations according to occupancy revealed that in cells of all sizes promoter/TF clustering narrowed the spread in average SBF residency time across all promoters, thus homogenizing SBF occupancy across promoters (Figure 3-Supplemental Figure 1C). This effect was particularly pronounced if the average SBF residency time was assessed for short periods (e.g., a 1 second test time) in large cells close to the G1/S transition (Figure 3-Supplemental Figure 1D). In this situation, clustering reduced the number of G1/S promoters that were never bound by an SBF complex during the test time by ∼2-fold. If the G1/S transition was triggered in this time window, the expression of all SBF-bound genes would be more correlated. This result suggests that clustering might facilitate the synchronous expression of the G1/S regulon.

### Scaling arguments explain transcription factor clustering

Scaling arguments provide a plausible explanation for why DNA-binding TF complexes (i.e., Swi4 and Mbp1 dimers) would be clustered at G1/S promoter clusters. Particle-DNA binding/unbinding is at equilibrium when the rate of binding events equals the rate of unbinding events. The TF binding/unbinding dynamics are in the 10-20 ms range (based on the observation of TF motion in live cells on the PALM frame timescale, see below) and on the detectable decrease of RICS vertical correlations, Figure 4-Supplemental Figure 3). Consistently, single particle tracking data with a spatial resolution of a few nanometers, i.e. much smaller than observed particle cluster size, showed a very minor fraction of completely immobile particles, buttressing the conclusion that particle dynamics is faster than the acquisition frame time of 30 ms. Thus, the rate of TF dissociation from DNA is of the order *k_off_*∼1⁄15ms=67s^-1^. Our previous model indicated that SBF/MBF dissociation constants are in the range of *K*_*S*3_ = 0.02 *μM*, corresponding then to a *k_on_* rate of:

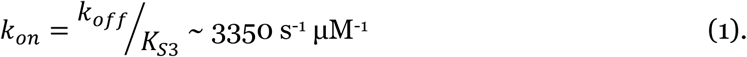

In the situation of a single TF dimer particle moving in a neighborhood of volume *V_o_* (in fL) containing *n* DNA binding sites evenly distributed, the propensity for this particle to bind DNA is:

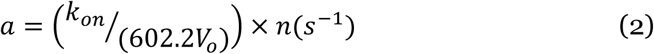

and therefore the mean free time (average time spent diffusing around without binding DNA) is:

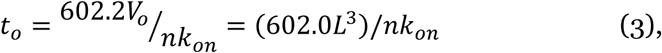

where *L* (μm) is the characteristic length defining the volume *V_o_*. During this time lag the particle diffuses away from its original point and jumps an average distance, *L_jump_*, of:

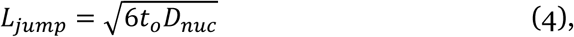

where *D_nuc_* = 2 − 3 *μm*^2^*s*^−1^. If DNA binding sites, rather than being clustered, are equally distributed within the nucleus, L∼1μm (nuclear size), *n*∼200 (number of G1/S promoters), and thus *t_o_*= 0.8-1 ms and *L_jump_*=100-200 nm. Thus, before binding to DNA again, the freely diffusing nuclear SBF/MBF particle explores a significant fraction (10-20%) of the nuclear radius and will therefore rebind at a location distant from its previous binding site. Thus, the diffusing particles are strongly mixed throughout the entire nucleus making random cluster formation unlikely (Figure 3-Supplemental Figure 2). Active processes may cluster the transcription factors to counteract diffusion, but we do not consider this possibility in the model, nor is it required for the model to achieve TF clustering.

Indeed, if on the other hand, DNA binding sites are pre-organized as clusters, *L*∼0.03 μm (cluster size), *n*∼6-8 (number of binding sites per cluster), then *t_o_*∼0.5-1 μs and *L_jump_*=3-5 nm, at most. Thus, *L_jump_*≪*L*, the cluster size. This implies that the next binding of the SBF/MBF particle will be within the same cluster of DNA sites where it was previously bound, and thus that diffusing SBF/MBF particles become dynamically trapped in G1/S promoter clusters. This trapping effect stems from a delicate balance between the density of DNA promoters within clusters and fast DNA-TF binding/unbinding dynamics that tend to favor clustering on one hand, and (fast) nuclear diffusion *D_nuc_* on the other hand that tends to dissolve clusters. In support of this argument, simulations with 100-fold slower binding/unbinding dynamics showed slightly reduced cluster number and/or size for all proteins (Table 1, compare rows 1 and 2).

**Table 1.** Sensitivity analysis of TF clustering simulations in very small cells (10.1fL).

| Parameters | Reaction | Number of Molecules/cluster |  |  | Number of Clusters/nucleus |  |  |
| --- | --- | --- | --- | --- | --- | --- | --- |
|  |  | Swi4 | Mbp1 | Swi6 | Swi4 | Mbp1 | Swi6 |
| $k_{on1,3\ s/m} = 3500$<br>$k_{on2,4\ s/m} = 35000$ | Default parameters | 4.27 | 5.15 | 5.38 | 2.2 | 10.1 | 15.2 |
| 100 fold lower<br>$k_{on1,2,3,4\ s/m}$ | All dynamics slow down | 4.09 | 5.26 | 5.38 | 2.1 | 9.7 | 13.9 |
| 20 fold larger $K_{s/m1}$ | With $k_{on1}$ constant this is an increased rate for Swi4d/Mbp1d dissociating from DNA relative to SBF/MBF | 4.27 | 4.90 | 5.39 | 2.2 | 11.8 | 17.0 |
| 20 fold larger $K_{s/m2}$ | With $k_{on2}$ constant this is an increased rate for Swi6 dissociation from Swi4/Mbp1 | 4.09 | 4.97 | 4.33 | 2.3 | 11.5 | 0.6 |

$k_{on1s/m}$ – On rate for Swi4/Mbp1 binding to DNA, $k_{on2s/m}$ – On rate for Swi6 binding to Swi4 or Mbp1, $k_{on3s/m}$ – On rate for SBF/ MBF binding to DNA, $k_{on4s/m}$ – On rate for Swi6 binding to Swi4/Mbp1 DNA complexes, $K_{s/m1}$ – Dissociation constant for monomer Swi4/Mbp1 binding to DNA, $K_{s/m2}$ – Dissociation constant for monomer Swi6 binding monomer Swi4 and Mbp1

In contrast, newly synthesized particles produced as cells grow populate new clusters rather than being trapped in existing clusters because of a simple saturation effect. If a diffusing particle is within the neighborhood of a partially occupied cluster, then the mean-free time becomes:

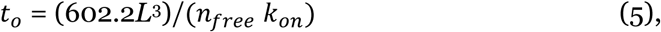

where *n_free_* is the number of free sites. If all sites are occupied then *t_o_*→∞, but even if there is one site free, the mean free time can increase by an order of magnitude, yielding a 3-4-fold increase in *L_jump_*, which becomes comparable to the cluster size. In this case, particles approaching nearly saturated clusters will have a strong probability to diffuse away without being captured, explaining why the number of particles trapped within each DNA cluster does not significantly increase with cell size even though SBF/MBF particle counts increase, and thus why the newly synthesized SBF/MBF factors tend to populate new DNA clusters. We have carried out the simulations using a 3-fold smaller (10 nm) and a 2-fold larger (60 nm) mesh size. Clustering was observed in both cases (Figure 3 – Supplemental Figure 3). Thus, the geometric representation of the clusters does not impact cluster formation in our model.

One of the surprising observations in both the experiments and the modeling was that TF clusters were observed in small cells in which the TF copy numbers are much lower than the number of target sites. These observations imply that the TFs do not distribute randomly to all promoters. (Note that in the model, all promoters had the same affinity). The mechanism described above can explain why Swi4 dimers/SBF and Mbp1 dimers/MBF particles would cluster naturally around pre-organized G1/S target promoter clusters. However, it does not explain why in small cells, where the total number of fully formed SBF/MBF complexes is 40-70, *i.e*. larger than the number of putative DNA clusters, only a small number of DNA clusters are populated by the TFs while many remain unoccupied.

To address this question, we focused on Swi6 clustering. Given the model parameters chosen and the species concentrations at equilibrium, the systems evolves in a regime where most Swi4/Mbp1 molecules are Swi6-bound, and hence promoters are mostly occupied by SBF/MBF (Figure 3-Supplemental Figure 1A-B). Swi6 does not directly bind to DNA: Swi6 clustering is dependent on its interaction with Swi4 and Mbp1. The dissociation of Swi6 from Swi4/Mbp1-bound DNA, which occurs regularly given the *k_off_* values, creates a Swi6-enriched region around a partially populated cluster, that can trap freely-diffusing Swi4 and Mbp1 dimers. This is not the case around empty clusters. This local concentration effect would favor the formation of new SBF/MBF complexes in the neighborhood of partially populated clusters, and thus the DNA binding of nearby Swi4d/Mbp1d dimers.

We computationally tested the hypothesis by modulating SBF/MBF affinity for DNA and Swi6 affinity for Mbp1/Swi4. In small cells, strengthening SBF/MBF DNA binding enhanced Swi6 clustering by increasing the molecules per cluster (Table 1, compare rows 1 and 3 for Swi6, last column). In contrast, decreasing the Swi6 affinity for Swi4 and Mbp1 markedly reduced the number and size of Swi6 clusters (Table 1, compare rows 1 and 4 for Swi6). Both these results directly follow from the fact that Swi6 forms clusters via binding to Swi4/Mbp1. Interestingly, decreasing the Swi6 affinity for Swi4 and Mbp1 also slightly reduced the molecular content of Swi4/Mbp1 clusters, while increasing cluster counts (Table 1, compare rows 1 and 4 for Swi4 and Mbp1), in agreement with a scenario where local Swi6 enrichment contributes to Swi4/Mbp1 accumulation at already populated promoter clusters.

Collectively, these local avidity effects offer one possible rationale for the observation of clusters for all three proteins even at low copy number in small cells. In this case, the number of molecules per cluster is set by the trade-off between two phenomena. The first is the saturation effect of target sites within each cluster as discussed above. Higher saturation within clusters increases the mean free time and the diffusion jumps between unbinding and re-binding events, favoring escape from clusters close to saturation and binding to empty promoter clusters. The second is the local Swi6-enrichment around partially populated clusters that improves the likelihood of binding new Swi4/Mbp1 molecules as full S(M)BF complexes to these clusters. We note however, that other factors such as chromatin states, other protein partners or different promoter affinities, operative in live cells, but not present in our model could also contribute to the observed clustering in small cells.

### Single particle tracking PALM in live cell nuclei reveals multiple modes of TF mobility

A key prediction of our mathematical model is that the G1/S TFs should display very different kinds of motion ranging from slow, confined diffusion within the neighborhood of promoter clusters to faster diffusion between clusters (Figure 4A, B, Figure 3-Supplemental Figure 4). In our model, this combination of slow and fast motion modes along the same single molecule trajectories, with relative weights that mirror the fractions of time spent within and between clusters, yielded effective diffusion coefficients ranging over orders of magnitude (Figure 4B, Figure 3 – Supplemental Figure 4C). This behavior was characteristic of anomalous sub-diffusion, exhibiting downward curvature of the Mean Squared Displacement (MSD) curves (Figure 3 – Supplemental Figure 4B).

**Figure 4.**
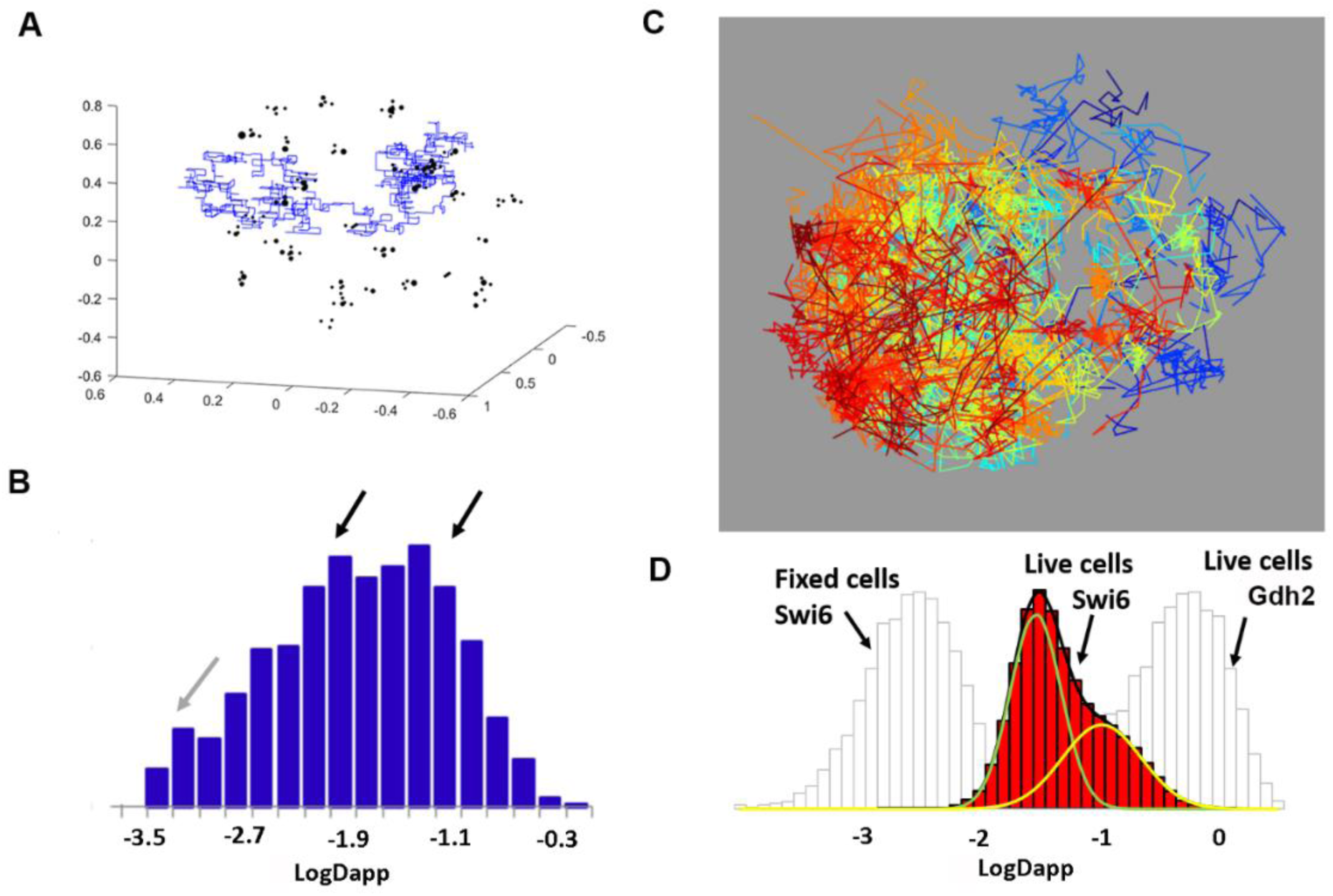
Swi4, Mbp1 and Swi6 dynamics are comparable in simulations and experiment. A) Example of individual molecule trajectory of Swi6 dimers from the simulations. B) Histograms of the log of the apparent diffusion coefficient of Swi6 from all simulation trajectories. Black arrows highlight two modes of motion. A third slower mode (grey arrow) likely arises from Swi6 dimers that spent most of their time bound to Mbp1/Swi4 DNA complexes during the 5s window of Monte Carlo simulations, and cannot be resolved in experiments due to instrument jitter and localization uncertainties. C) Experimental individual molecule trajectories of Swi6-mEos3.2 in the nucleus of a live cell grown in SC+2%glucose medium. Each individual trajectory is represented by a different color. D) Histogram of the log of apparent diffusion coefficients, Dapp, calculated from all individual trajectories of Swi6-mEos3.2 for all cells in three FOV (red) (∼5000 trajectories per FOV). The distribution was fit to two Gaussians. The distribution for Swi6 Log Dapp obtained from spt tracking in fixed cells (light grey - immobile, except for instrument jitter) and the distribution of Log Dapp of mEos3.2-Gdh2 (glutamate dehydrogenase)-mEos3.2 (also light grey) diffusing freely in the cytoplasm are plotted for comparison and indicated by the arrows.

To evaluate the dynamic properties of our model, we measured the dynamics of Swi4, Swi6 and Mbp1 using single particle tracking (spt)PALM in live cells (Videos 1-3). After analysis of the trajectories (see Methods), images of overlaid individual trajectories for each nucleus were produced (Figure 4C). We note the similarity of the image in Figure 4C, in terms of the space mapped out by the overlaid trajectories with previous models of the yeast nucleus (Duan et al., 2010; Wong et al., 2012). The trajectories of individual molecules were a mixture of smaller and larger MSDs (Figure 4 – Supplemental Figure 2A, B). Analysis of the individual experimental MSD in terms of apparent diffusion coefficients at short timescales yielded effective (single molecule) diffusion coefficients whose distribution could be fitted reasonably well with superpositions of two Gaussian modes (Figure 4D, Figure 4 - Supplemental Figure 2C). Experimental trajectory length distributions were similar in both carbon sources tested, and the majority were shorter than 25 frames (Figure 4-Supplemental Figure 1).

The average apparent fast diffusion coefficients for each TF were approximately 0.1 μm^2^/s in both simulations and experiments (compare Figures 4B rightmost arrow, 4D yellow gaussian, and Figure 3 – Supplemental Figure 4, Figure 4 – Supplemental Figure 2), ∼10-20-fold slower than that observed for glutamate dehydrogenase (mEos3.2-Gdh2), which diffuses freely in the cytoplasm. This value was also considerably slower than the diffusion coefficient of free nuclear proteins evaluated by Raster Scanning Image Correlation Spectroscopy (RICS) (Thattikota et al., 2018). The lower mobility of this component, with respect to free protein diffusion, is consistent with the fixed cell data showing that 10-15% of the molecules are outside of the clusters and are likely undergoing a combination of free diffusion and non-specific DNA binding. The much slower apparent diffusion coefficient, 0.01-0.03 μm^2^/s, was consistent with confinement of ∼85% of the TFs in clusters. This motion was 10-fold faster than the apparent diffusion of immobile molecules in fixed cells (which arises from instrument jitter and localization imprecision) (Figure 4D). This slow TF diffusion component of 0.01-0,03 μm^2^/s constitutes a signature of actual TF motion on the scale of the PALM frame time (30 ms), supporting the values of <30ms for the off-rates of DNA-TF complexes (*k_off_*) used in our modeling. We note that the apparent diffusion coefficient of this pool of slowly moving G1/S TFs is also at least one order of magnitude faster than the diffusion coefficient previously reported for chromosomal motion (estimated to about 0.0005 μm^2^/s, (Marshall et al., 1997)), although faster chromosomal motion could occur on the PALM frame time. Arbitrary Region Raster Scanning Image Correlation Spectroscopy (ARICS) analysis (Schrimpf et al., 2018) of Swi4-, Mbp1- and Swi6-GFP fluorescence fluctuations for nuclear pixels only yielded diffusion coefficients of 0.015, 0.05 and 0.11, respectively for the three proteins (Figure 4 – Supplemental Figure 3), also in reasonable agreement with the experimental sptPALM results and simulations.

Since individual molecules sampled both fast and slow dynamic modes in a single trajectory, these apparent diffusion coefficients conflate the two modes of motion, which results in the apparent anomalous nature of TF diffusion. To avoid mixing different modes of motion over single trajectories, we analyzed the dynamics of individual molecules using a Jump-Distance Distribution (JDD) approach (Menssen and Mani, 2018; Tollis, 2015). JDDs for Swi4, Swi6 and Mbp1 in glucose and glycerol-grown cells (Figure 4-Supplemental Figure 4B bottom) all showed a main peak (black arrows) corresponding to a Jump Distance ranging from 30-50 nm (for Swi4) to 60-80 nm (for Swi6 and Mbp1), for a duration of 7*30 ms=0.21 s (8 trajectory points), in agreement with apparent cluster sizes on PALM images. Furthermore, the position of this peak was barely altered when plotting the JDD on larger timescales (18 trajectory points, 17*30ms=0.51s, Figure 4-Supplemental Figure 4B top), emphasizing that this peak corresponds to confined molecules, for which the volume of diffusion (and hence the jump distance) does not depend on the sampling time. In comparison, free nuclear diffusion in our quasi-2D excitation volume would yield a typical jump of ∼1 µm, emphasizing how strongly the molecular motion is restricted. Comparison of JDDs acquired in live (Figure 4-Supplemental Figure 4B) and fixed (Figure 4-Supplemental Figure 4A) cells revealed that jump distances are significantly larger in live cells, indicating that the peak at small jump distance in this case is not due to instrument jitter but represents actual motion. The long tail in the JDD in live cells (Figure 4-Supplemental Figure 4B, red arrows) indicated that a small fraction of the particles displays fast motion, in agreement with the MSD analysis and model predictions.

The JDDs could not be well fitted using simple models such as free diffusion, anomalous diffusion, anisotropic motion along (and around) linear tracks or with a mixed two-mode model of free diffusion. Rather, our live cell data were best characterized by a superposition of an anomalous diffusion component (low JDD main peak, 65-80% of the molecules, see Table 2 columns 3-4) with a faster anisotropic motion (20-35% of the molecules) characterized in our fitting by a unidirectional motion along a linear track (Table 2, column 5) with variance around the track (Table 2, columns 6-7). The latter component may correspond in part to anisotropic diffusion along DNA (Von Hippel and Berg, 1989).

**Table 2.** Jump Distance Distribution parameters from live cell spt-PALM experiments.

| | Fraction of anomalously diffusing molecules (%) | Anomalous diffusion coefficient ($\mu\text{m}^2/\text{s}^a$) | Anomalous power (a) | Velocity along tracks ($\mu\text{m}/\text{s}$) | Variance around tracks | Diffusion coefficient around tracks ($\mu\text{m}^2/\text{s}$) |
| --- | --- | --- | --- | --- | --- | --- |
| Mbp1, glu | 66.26 | 0.0091 | 0.73 | 0.1198 | 0.007833 | 0.1263 |
| Mbp1, gly | 68.80 | 0.0080 | 0.74 | 0.1043 | 0.00760 | 0.1226 |
| Swi4, glu | 80.80 | 0.0054 | <0.4833 | 0.1577 | 0.006205 | 0.1001 |
| Swi4, gly | 81.64 | 0.0044 | 0.6038 | 0.5251 | 0.004683 | 0.0755 |
| Swi6, glu | 65.46 | 0.0089 | 0.8263 | 0.0976 | 0.007743 | 0.1249 |
| Swi6, gly | 75.56 | 0.0065 | 0.6767 | 0.1066 | 0.006898 | 0.1113 |

These results were also in good quantitative agreement with the cluster analysis in fixed cells that yielded ∼15% and ∼85% of TFs outside and inside clusters respectively at the time of fixation. Thus, at the short timescale of JDD computation, TFs exhibited anomalous motion, confirming that DNA-binding/unbinding dynamics were faster than the ∼100ms regime. We note that both JDD and MSD analyses were performed across time intervals that exceeded 100ms, a time interval during which fast diffusing molecules (D∼2-3μm^2^/s) travel an average of 400-500nm. This (linear) distance is comparable to the depth of our microscope field in the z-direction, and therefore a small fraction of such fast diffusing molecules might escape the microscope field and not be accounted for in our analysis. Thus, it is possible that we slightly underestimate the fraction of fast-diffusing molecules. Analysis using either the JDD or the two component MSD of complete trajectories revealed that Swi4 is much less mobile than either Swi6 or Mbp1, perhaps be due to higher affinity (lower off-rates) of Swi4 for its target sites on DNA. This was also the case for the ARICS analysis above. Moreover, Mbp1 and Swi6 mobility decreased in glycerol medium compared to glucose (Figure 4-Supplemental Figure 2). This result is consistent with stronger specific binding of Mbp1 in the poor carbon source and is in agreement with the conditional large cell size phenotype of the *Δmbp1* mutant reported previously (Dorsey et al., 2018). Collectively, these results suggest a mechanism whereby G1/S TFs populate discrete clusters and can also jump between clusters.

## Discussion

Super-resolution spatial mapping of Swi4, Mbp1 and Swi6 molecules in fixed cells revealed that these TFs do not distribute randomly but are organized into discrete clusters of ∼8 molecules, even in small cells, and that the number of clusters increases as cells grow. Stochastic modeling suggests that the spatial organization of the G1/S TFs is linked to the underlying spatial organization of their ∼200 target promoters. The distribution of G1/S TFs cluster sizes (in number of molecules) that we observed strongly resembles the multinomial distribution of promoter cluster sizes that one would expect by randomly distributing ∼200 promoters across 30-35 clusters (i.e., typically 4-12 promoters per cluster), although we cannot exclude that larger clusters are formed *in vivo* via biologically active processes. Although our results do not rule out explicitly the converse possibility that G1/S TFs might spontaneously assemble into clusters, any such mechanism would need to counteract free diffusion. Spontaneous assembly would also favor aggregation into a one or a few large clusters of variable size, rather than the size dependent accumulation of discrete clusters that we observe.

Importantly, clusters are observed in small cells where the TFs are severely limiting with respect to promoter target sites, and the number of molecules per cluster does not change with increasing copy number or size. Our simulations demonstrate that the balance between Swi6 local concentration effects on the one hand and target site saturation versus diffusion propensity on the other, is sufficient to explain the existence of clusters in small cells and the observed successive titration of new clusters as cells grow. The simulations suggest that the occupation of some sites within a cluster by SBF or MBF tends to sequester Mbp1 and Swi4 molecules via transient interactions with the Swi6 activator already present in the cluster. However, as target sites within any given cluster are bound by the increasing TF copy numbers as cells grow, the decreased number of unbound target sites available decreases the Swi4 or Mbp1 binding propensity. Diffusion out of the cluster eventually becomes statistically more probable than DNA re-binding. This interpretation is consistent with the bimodal dynamics of the G1/S TF we observe by sptPALM in live cells. Overall, these results indicate that cluster size and the distribution of TFs across clusters can be tuned by the promoter content of pre-formed clusters and by relative affinities of TF subunits for each other and for target sites on DNA. We note that these modeling results, based on known concentrations and reasonable affinity and diffusion constants, account well for the observation of clusters even in small cells, but do not rule out other mechanisms of cluster formation.

Both general and specific transcription factors have been observed to form clusters. For example, in budding yeast, the transcriptional repressor Mig1 forms clusters of similar size as we observe for Swi4, Mbp1 and Swi6 (Wollman et al., 2017). Interestingly, the number and copy number content of Mig1 clusters increases upon glucose repression, and like the G1/S TFs, these clusters also exhibit mixed dynamic properties (Wollman et al., 2017). In mammalian cells, RNA Polymerase II and its associated Mediator complex are co-localized in much larger stable clusters (>300 nm, ∼300 molecules) that exhibit properties of phase separated condensates (Cho et al., 2018). Even larger clusters have been observed using super-resolution imaging for the transcription factor STAT3 (Gao et al., 2017). These examples suggest that TF clustering is a common phenomenon, but that different TFs can exhibit different clustering behavior.

While our observations can be accounted for by simple physical phenomena and captured in a mathematical model based on minimal assumptions, the observed clustering of the G1/S TFs must be coupled at some level to the global organization of the yeast genome (Duan et al., 2010; Lazar-Stefanita et al., 2017; Taddei and Gasser, 2012; Wong et al., 2012; Zimmer and Fabre, 2011). Thus, our results lead to several open questions. The nature of the pre-formed G1/S promoter clusters inferred from our model is unknown at this juncture, but may be generated by condensin- and cohesin-mediated chromosome looping (Lazar-Stefanita et al., 2017), perhaps in conjunction with other factors that bind specific promoter regions or some other feature of global genome organization. Self-associating domains of 1-5 genes, with boundaries at highly transcribed genes, have been reported in yeast chromatin (Hsieh et al., 2015), although they are smaller than the Topologically Associated Domains (TADs) typically found in mammalian cells (Bintu et al., 2018; Cattoni et al., 2017; Dixon et al., 2012; Nora et al., 2012; Sexton et al., 2012; Szabo et al., 2018). In quiescent cells self-associating domains in yeast are in a repressed state, apparently organized by condensins (Swygert et al., 2019).

It remains to be determined if the clusters observed here are populated in a discrete order as cell grow and whether clusters have defined or random promoter compositions. It is also unclear whether clusters can exchange promoters over time as opposed to being of fixed composition. Regardless of the underlying static or dynamic mechanisms, the localization of G1/S promoters within discrete clusters may help coordinate the G1/S transcriptional program once Start is triggered. These results provide evidence that higher-level organization of the genome may contribute to the efficiency of cell state transitions that depend on complex gene regulons.

## Methods

### Key resources table

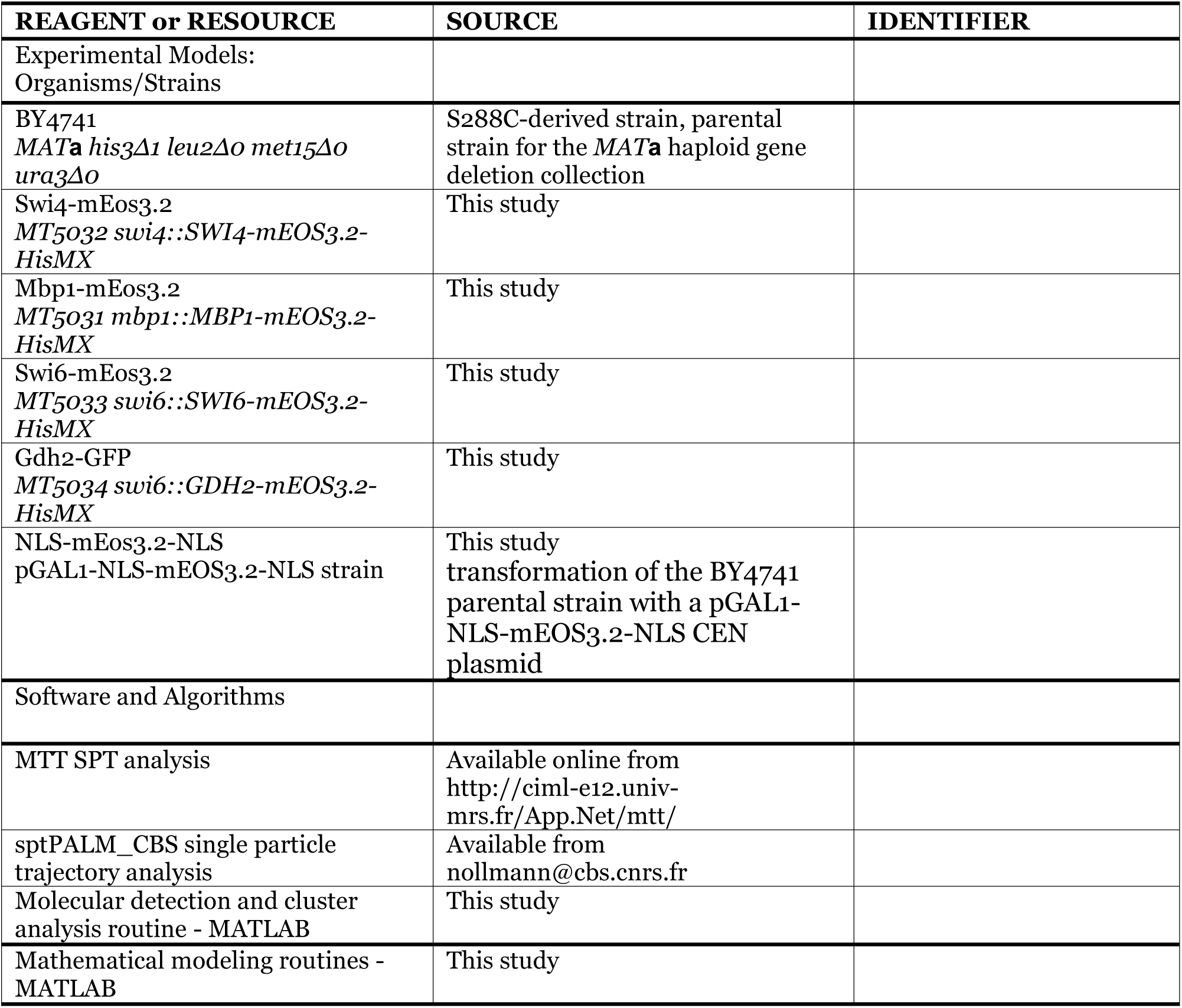

### Strains and sample preparation

The *S. cerevisiae* strains used in this study were constructed in the S288C (BY4741) background by PCR-based homologous recombination integration of a mEOS3.2-HisMX cassette at the 3’ end of each reading frame at the endogenous loci. The strain genotypes were:

BY4741 MAT**a** his3Δ1 *leu2Δ0 met15Δ0 ura3Δ0*

MT5031 *mbp1::MBP1-mEOS3.2-HisMX*

MT5032 *swi4::SWI4-mEOS3.2-HisMX*

MT5033 *swi6::SWI6-mEOS3.2-HisMX*

MT5033 *swi6::GDH2-mEOS3.2-HisMX*

The pGAL1-NLS-mEOS3.2-NLS strain was obtained by transformation of the BY4741 parental strain with a pGAL1-NLS-mEOS3.2-NLS CEN plasmid that was constructed by Gibson assembly of a SV40 (NLS) domain (Gblock, IDTDNA), the mEOS3.2 sequence amplified from our mEOS3.2-HisMX cassette, and another SV40 (NLS) domain (Gblock, IDTDNA) into a pGAL1 CEN plasmid.

The TF-mEos3.2 fusion strains were grown on synthetic complete (SC) –His + 2% glucose plates for 3 days. Fresh colonies were picked and grown in SC–His media supplemented with a rich (2% glucose) or poor (2% glycerol) carbon nutrient source until stationary phase. Prior to imaging, strains grown on rich or poor carbon sources where diluted to 0.3 OD and allowed to grow until OD 0.7 in fresh SC + 2% glucose/glycerol (His-) medium. A 1 mL sample of OD 0.7 culture was pelleted at 3000 rpm and washed with fresh media. The sample was concentrated 10x by removing 900 mL of media and the cells were resuspended. A 5 µL sample of the cells was placed on a concanavalin A (ConA) coated #1 coverslip with 100 nm Tetraspec fluorescent beads and allowed to adhere to the surface for 4 min. The cover slip was then placed on a 2% agar SC – (His-) + carbon source pad as previously described (Dorsey et al., 2018) and immediately imaged. For fixed cells, a 1 mL sample of cells was washed with PBS buffer, buffer was removed and then 500 µL of a 4% paraformaldehyde solution was added and allowed to react for 20 min. The sample was washed extensively with PBS and place on a #1 cover slip coated with ConA and 100 nm Tetraspec fluorescent beads similar to live cells. Fixed cell samples were treated the same as live cells from this point on.

### PALM Microscope

Imaging was performed with a custom-built PALM/Storm system based on a Nikon inverted Ti-U Eclipse microscope and similar to that described in (Fiche et al., 2013). A CFI Plan Apo Lambda 100x/1.40 NA Oil Objective was used and images were collected on an Andor iXon Ultra 897 EMCCD camera. A 561 nm laser (Sapphire 561-150 CW CDRH) at 0.2 kW/cm^2^ and a 405 nm laser (OBIS 405 nm LX 50 mW) at 0.3 W/cm^2^ were directed into the microscope objective in a Koehler illumination configuration with the aid of a pair of beam-expanding lenses (150/30 mm, Thor Labs) and a quad-band (405/488/561/640 nm, Chroma) filter for mEos3.2 excitation and activation, respectively. Emission was collected with a 600/50 nm BP filter (Chroma) mounted in an automated filter wheel (Thor Labs). The image was magnified with a set of lenses (150/250 mm) to an effective pixel size of 120 nm creating a 46×46 µm^2^ imaging area. Excitation and activation power were controlled using an acousto-optical tunable filter (AOTFnC400.650-TN, AA Optoelectrionics). The imaging focal plane was locked in position (within 10 nm) using an autofocus program by tracking the reflection of a 785 nm IR source (OBIS 785 nm LX 100 mW) from the sample coverslip with a Thor Labs CCD camera and continuous incremental adjustments with a Fast PIFOC Piezo Nanofocusing Z-Drive (PI) (Figure 5C) (Fiche et al., 2013). Software controls and data acquisition for the microscope stage, laser excitation and activation power, autofocus and camera were written in Labview 2015 (National Instruments).

**Figure 5.**
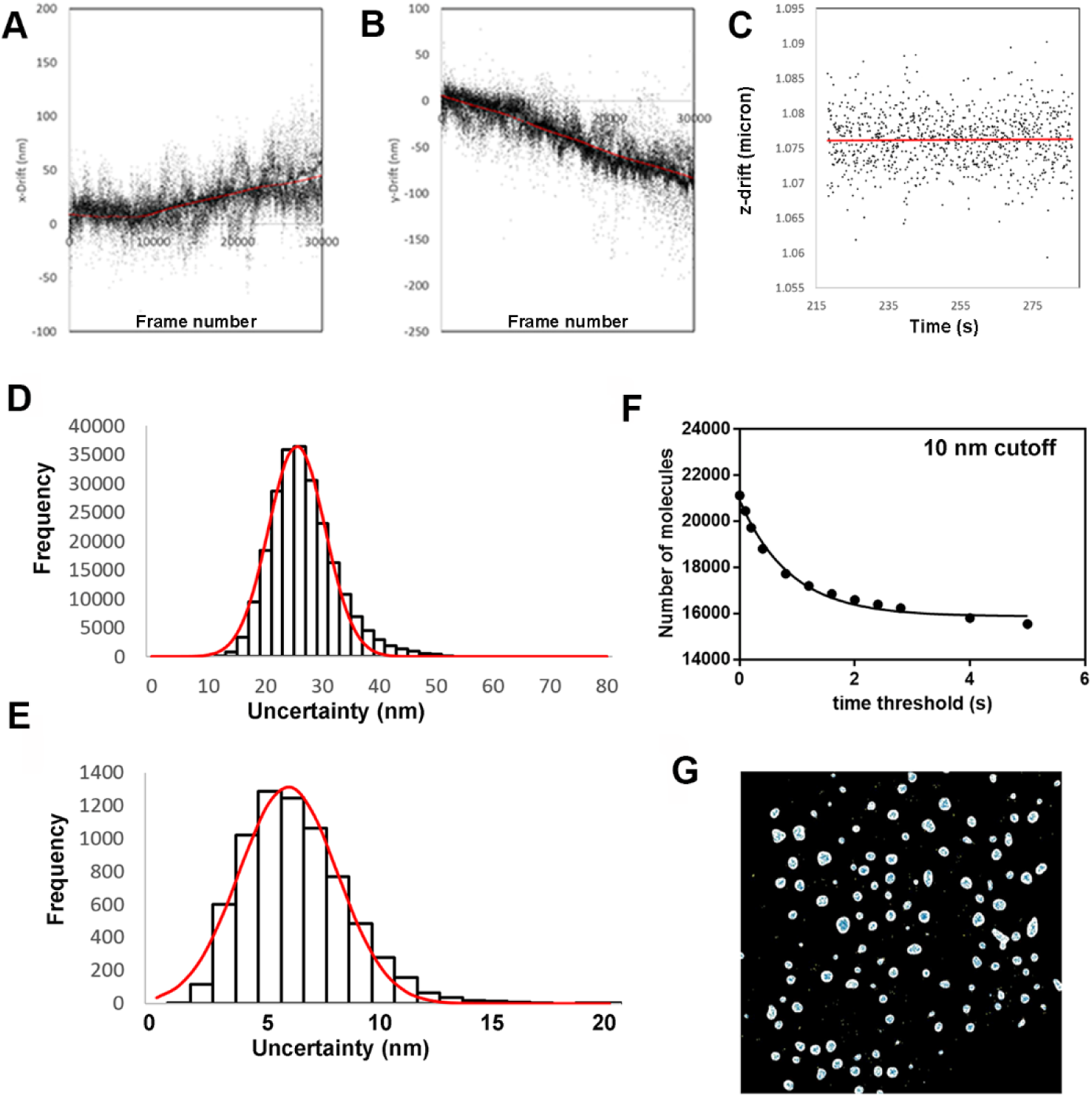
PALM Acquisition and analysis. A-C) Positional drift correction in x, y and z-axes respectively, as described in the Methods section and SI. D) Detection localization uncertainty histogram for all detections in a typical FOV. E) Standard error on the mean of the molecular localizations for a typical FOV. F) Blinking time constant determination. G) Example of the results of the nuclear masking algorithm.

### Super-resolution image acquisition

Samples were placed in a cylindrical sample chamber and mounted on the microscope. After locking in the optimum imaging plane of the sample, a bright field reference image was acquired. For each FOV, 30,000-40,000 frames (exposure time 30 ms) were collected within a data acquisition. The 561 nm CW laser at a fixed power was used for excitation of mEos3.2 molecules in the red-shifted state. Photoactivation was achieved by continually increasing the power of a 405 nm CW laser diode until mEos3.2 photoswitching was no longer observed (Lee et al., 2012). For each FOV, 30,000-40,000 frames were collected for each acquisition. Active correction for drift in the z-plane was carried out as previously described (Fiche et al., 2013) (Figure 5A-C).

### PALM Analysis

PALM image stacks were analyzed and x-y drift-corrected using the Thunderstorm ImageJ plugin (Figure 5A-C). If images drifted more than 2 pixels (120 nm effective pixel size) in either x or y directions, then the data were discarded. For acquisitions within the threshold drift range, molecular detection positions were drift-corrected using the image-correlation drift algorithm from Thunderstorm that cross-correlated the signal from 100 nm fluorescent beads (Tetraspek, Anaspec) used as fiducial markers. The x-y drift was fit to a smoothing function with the frame number as the independent variable (Figure 5A, B). The average frame number for an individual molecule was then used to compute its total associated drift, which was added to its averaged x-y position.

The average resolution for each individual localization was 25 nm (Figure 5D), which allowed an optimal super resolution image pixel size of 12 nm. The MATLAB script MTT (Sergé et al., 2008) was used to connect molecular trajectories (i.e., positions of the same molecule from one frame to the next) and the sptPALM_CBS (Fiche et al., 2013) script was used to analyze and filter the trajectories. For fixed cell analysis, nuclei were masked with the MATLAB gaussfit function (Figure 5G) using our custom SuperResolution MATLAB script. This script also corrected for over-counting due to mEos3.2 blinking (Figure 5F) (Lee et al., 2012), with spatial and temporal cutoffs of 10 nm and 2 seconds. Since most molecules were found within the nucleus, the nuclear mask sets boundaries within which the number of molecules per nucleus, number of clusters per nucleus and number of molecules per cluster were determined. This procedure yielded the overlaid corrected detection images (see Figure 1A-DC, Figure 1-Supplemental Figures 1A-D and 2A-D). The distributions of masked nuclear sizes (Figure 2-Supplemental Figure 1) resembled the cell size distribution of asynchronous cultures, and therefore we used the mode of this distribution (∼20.000 pixels) to characterize the upper limit size of the G1 cells nuclei. Of note, in a spherical model for both cells and nuclei with a karyoplasmic ratio of 1/7 (Jorgensen et al., 2007), this size is equivalent to ∼30fL, a good estimate of the G1/S critical size of wild-type cells in SC+2%glucose.

The drift-corrected positions of each individual molecule detected in multiple (5-100) successive frames were averaged to yield the molecular images (see Figure 1E-G, Figure 1 - Supplemental Figures 1E-G, 2E-G and 3B, D, F). The pixel size in these images was chosen as 3 nm, half the value of the standard error on the mean for all molecular positions within a data set (6 nm) (Figure 5E). The number of molecules per nucleus and the cell size in pixels were also generated by this script. Live cell single particle tracking (spt)-PALM trajectories were analyzed as Mean Square Displacement (MSD) as a function of time shift along each trajectory using the sptPALM_CBS MATLAB script (Fiche et al., 2013). To disentangle distinct molecular motion modes along individual trajectories, trajectories were also analyzed using the Jump Distance Distribution approach (Menssen and Mani, 2018; Tollis, 2015) in which trajectories are subdivided into short sections (8 points, 210ms time bins) and analyzed collectively. Separate experiments were carried out on six different days for Mbp1, and three different days for Swi4 and Swi6 for a total of 28 FOV for fixed cells and 14 for live cells for Mbp1, 10 FOV for fixed cells and 6 FOV for live cells for Swi4 and 15 FOV for fixed cells and 8 FOV for live cells for Swi6. Despite the inherent limitations of PALM for particle counting, the super-resolution images of Swi4, Mbp1 and Swi6 correspond to a reasonable representation of their actual distribution in the nucleus for several reasons. First, non-activated (and hence unobserved) mEos3.2 molecules should be randomly distributed and hence unlikely to bias the fraction of molecules detected in clusters vs non-clustered molecules. Secondly, our low intensity, continuous switching illumination parameters allow for relatively efficient photoactivation. Moreover, because the three TFs diffuse as dimers (Dorsey et al., 2018), the majority of dimers are detected via at least one of their constituent monomers. Finally, yeast chromosomes bearing the G1/S TF target sites occupy a limited volume of the nucleus which limits problems of detection due to depth of field. Thus, the number of clusters per cell and the cell size invariance of cluster size are reasonably well determined.

### Fixed cells: cluster analysis

We used the nuclear super-resolution molecular images to develop custom cluster analysis scripts in the MATLAB environment. For each nucleus, we ordered the particles in a list such that each particle was next to its nearest neighbor in the nucleus using an algorithm based on the OPTICS ranking algorithm (Kriegel et al., 2011). This list is referred to as the nearest-neighbor ranked particle list. We computed the list of distances between each particle and the next across this list (see Figure 2-Supplemental Figure 3 and Figure 2-Supplemental Figure 5). Plotting this list revealed two characteristic features: valleys (red arrows), wherein the distance between consecutive particles is small, separated by spikes corresponding to large distances (blue asterisks). Valleys represent particle clusters, wherein the inter-particle distance is small, whereas spikes are characteristic of the distance between clusters. These plots provided a tool to count the number of clusters in the nucleus, through the definition of a distance threshold (red line in Figure 2-Supplemental Figure 3A), such that particles separated by a distance lower that the threshold were assigned to the same cluster. We computed the relative variation of the total number of clusters (across our entire dataset for each protein and each FOV, Swi4, Mbp1 and Swi6) as a function of threshold (see Figure 2-Supplemental Figure 3B). We found that the number of detected clusters is threshold-dependent for values lower than 10 super-resolution pixels (30 nm), whereas for values > 30 nm there is little dependence on the threshold. This indicates that most clustered particles were within a 30 nm neighborhood of their closest neighbor, a distance which thus represented a logical cluster size detection threshold. The nearest-neighbor ranked list of particles provided a simple means to count clusters within each nucleus, to count the number of particles within each cluster, and to correlate these data with nuclear size.

### Live cells: Mean square displacement and jump-distance distribution analyses

To analyze single particle tracking data and gain insight on the dynamic motion features at the molecular scale, we used two methods, each with complementary advantages and disadvantages. First, we selected entire individual trajectories, and computed the Mean Square Displacement (MSD) as a function of time shift along each trajectory (Figure 4-Supplemental Figure 2) using the sptPALM_CBS MATLAB script (Fiche et al., 2013). Although this analysis revealed a sub-diffusion type motion with confinement at large times, below 200ms the MSD curves could be fit reasonably well with a linear function. The slope provided the trajectory-averaged instantaneous diffusion coefficient (as shown in Figure 4, and Figure 4-Supplemental Figure 2). We processed simulated *in silico* individual trajectories in the same manner (Figure 3-Supplemental Figure 4).

Although MSD analysis is a powerful method to reveal particle confinement, because dynamics are averaged over entire trajectories (and further averaged across different trajectories), this approach only provided a global view of particle dynamics on the seconds timescale. To disentangle distinct molecular motion modes along individual trajectories, trajectories were also analyzed using the Jump Distance Distribution (Figure 4-Supplemental Figure 4) approach introduced in (Tollis, 2015) and further developed in (Menssen and Mani, 2018). Trajectories were subdivided into short sections (8 points, 210ms time bins) and analyzed collectively. From these sub-trajectories, a jump distance distribution (JDD, i.e., the distance covered along any given sub-trajectory within the 210ms of its duration) was computed for all data from a given experiment. This approach increases the likelihood of observing a unique mode of molecular motion along a shorter fraction of single molecule trajectories. We fitted the experimental JDDs with molecular motion models, including free (Brownian) diffusion, anomalous diffusion, directed transport along linear tracks, or more complex models that incorporates two of the classical motion models discussed above (i.e., where the population of sub-trajectories includes two subpopulations with different underlying transport modes). To select among these competing models, we used a Bayesian model selection procedure (see (Menssen and Mani, 2018; Tollis, 2015)), which outputs the probability of each underlying motion model in a manner that balances the fitting quality of a given model to its complexity.

### Mathematical modeling and Monte Carlo simulations

To model SBF/MBF formation and binding to target promoters, we used our previously published Start model (Dorsey et al., 2018). The model comprises a mass-action kinetics-based SBF/MBF binding module, resolution of which yields the concentrations of DNA-bound and DNA-free SBF/MBF complexes, and of Swi6-free Swi4/Mbp1 dimers bound to DNA in the cell nucleus as a function of cell size. From previous fluctuation microscopy-based measurements (Dorsey et al., 2018), we used the size-independent nuclear Mbp1 and Swi6 concentrations (respectively, 110nM and 150nM) and the Swi4 concentration that doubles linearly between early G1 (50nM in 14 fL cells) and late G1 phase (100nM in 35 fL cells). We assumed that interaction K_d_ values for the DNA binding proteins (Swi4 or Mbp1 = DBP) to DNA were unaffected by Swi6 binding, and vice-versa. In addition, our previous brightness data revealed that all measured Start proteins were predominantly dimeric (Dorsey et al., 2018). Thus, we reduced the model complexity by neglecting the equilibrium concentrations of protein complexes formed with monomer DNA-binding protein (DBP, Swi4 or Mbp1) and/or Swi6. As a result, in the steady state the SBF/MBF binding module reduces to 8 reactions (with effective dissociation constants *K_d_* derived in (Dorsey et al., 2018):

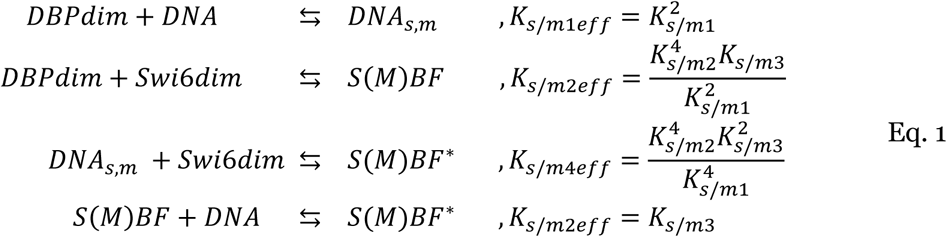

where *dim* stands for dimer and *DBP* can be either Swi4 or Mbp1, *DNA* and *DNA_s,m_* represent a DBP-free and DBP dimer-bound target promoter respectively, *S(M)BF* and *S(M)BF* ^∗^ are fully formed DNA-free and DNA-bound DBP dimer-Swi6 dimer SBF and MBF complexes, and the microscopic dissociation constants *K*_*s/m*1−3_ respectively characterize monomer DBP-DNA, monomer DBP-Swi6 and dimer DBP-DNA binding, with s and m lowercase subscripts standing for Swi4 and Mbp1, respectively. It is noteworthy that the dissociation constant of *S(M)BF* is not the microscopic DBP/monomeric-Swi6 constant but an effective dimer-DBP/dimer-Swi6 dissociation constant that involves multiple interactions. Unless otherwise specified, we used the following default values: *K*_*s*1_ = *K*_*m*1_ = 100*nM*, *K*_*s*3_ = *K*_*m*3_ = 20*nM*, *K*_*s*2_ = 20*nM*’ < *K*_*m*2_ = 50*nM* (Dorsey et al., 2018).

The mass action-like ordinary differential equations corresponding to this equilibrium relate the variation of the concentrations of the transcription complexes (right-hand side of Eq.1) to their formation rates (*k*_*on*1−4*s/m*_ *[interacting species], left-hand side of Eq.1) and their dissociation rates (*k*_*off*1−4*s/m*_ = *K*_*s/m*1−4*eff*_ ∗ *k*_*on*1−4*s/m*_ x [TF complex]):

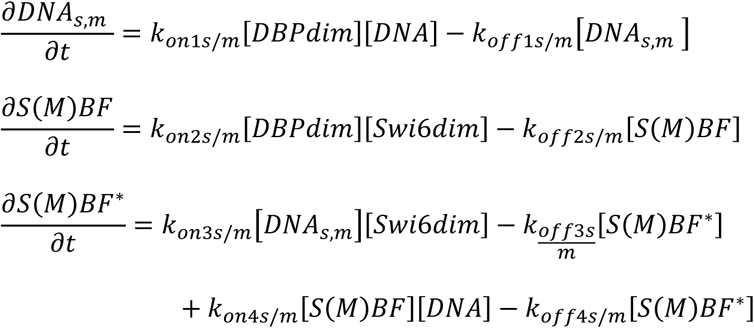

Default kinetic on-off rates are given in main text Table 1. These ODEs were converted to stochastic simulations using the Gillespie algorithm modified to account for diffusion (David Bernstein, 2005; Jose et al., 2013). The Gillespie algorithm is a particular class of Monte Carlo simulation algorithm originally developed to stochastically simulate biochemical systems with molecules binding to and dissociating from each other in a homogeneous, well-mixed solution (Gillespie, 1976). For any given state of the system at a given time, the algorithm associates each type of species (individual molecule or complex) a propensity (in s^−1^) to convert into another (i.e., a reaction). The sum of all propensities is then used to randomly determine the time to the next reaction, and another random number is generated to determine the type of the next reaction such that reactions with high propensities are more likely to be chosen than reactions with low propensities. Propensities are next re-evaluated at the new time, and successive iterations of this algorithm simulate the stochastic behavior of the chemical system as a function of time. The Gillespie framework has been successfully used by us and others to address various cell biological questions including, for instance, RNA secondary structure folding kinetics (Clote and Bayegan, 2018), stochastic gene expression (Ferguson et al., 2012) and yeast polarity establishment (Jose et al., 2013).

The inclusion of diffusion in this framework is straightforward. It requires partition of the reaction-diffusion volume into small elements. It considers identical molecules in different elements as different diffusing-reacting species. Finally, it extends the list of possible reactions between species, such that regular reactions (converting species A into B) are only possible between molecules within the same volume-element i (A,i -> B,i), and that molecule A diffusing from one element i to a neighbor j is a reaction that converts the species A,i into the species A,j. We divided the nuclear volume into infinitesimal volume elements (in 3D Cartesian coordinates, with xyz mesh-size =30nm, the maximal size that still provides sub-cluster resolution). We defined for each diffusible particle (i.e., free Swi4, Swi6, Mbp1 dimers, and DNA-free SBF and MBF): 1) a propensity to isotropically diffuse to a neighboring element (in 6 directions, see grey arrows), 2) bind to another particle (black arrows pointing towards each other) or 3) bind to DNA (black arrows pointing towards a DNA site), as defined by the model Eq.1 (see Figure 3A). In contrast, DNA-bound species are not assumed to diffuse at all, since chromosomal motion is expected to be negligible on the short time scale of our measurements and simulations (<2s, 10s respectively) (Marshall et al., 1997). However, DNA-bound species have propensities to dissociate from each other or from DNA (black arrows pointing away from DNA sites) according to the model Eq.1. Importantly, for a given set of on rates and microscopic K_d_’s (which define the effective K_d_’s for dimer interactions, see (Dorsey et al., 2018)) propensities depend on the mesh size *h* (Figure 3A). This follows from the fact that propensities are defined for individual particles or pairs of particles for complex formation. In the latter case, for any individual particles of type A and B, the rate of A-B complex formation depends linearly on the apparent concentration of particle B, which is proportional to the inverse of the element volume 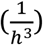. This term is absent in complex dissociation events (David Bernstein, 2005). The presence of the term 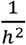 in the propensity of diffusion events follows from the second order spatial derivative in the diffusion equation. One critical condition for the use of the Gillespie algorithm is that the reaction-diffusion volumes are well mixed. This holds as long as the simulated number of diffusion events is significantly larger than the number of biochemical reaction events. This condition is fulfilled in our simulations since diffusion events typically exceed reactions by 2-3 orders of magnitude.

This algorithm was implemented numerically in MATLAB, for small, medium and large cells with nuclear radii of 0.67, 0.8 and 0.98 µm (corresponding to cell volumes respectively of 10, 17.1 and 31.5 fL), using a cell size-independent karyoplasmic ratio (Jorgensen et al., 2007) on a h=0.03 µm three-dimensional mesh and 200 DNA promoters randomly distributed across 35 clusters, themselves randomly distributed within the nucleus for each simulation. Specifically, each promoter assigned to a cluster was either positioned within the element containing the cluster center, or in an immediate neighbor element (a total of 7 possible positions). Given the mesh size of h=30nm, this procedure ensured that DNA promoters belonging to the same cluster are within a 30-60nm distance from each other, in agreement with observed cluster sizes (Figures 1E-G, Figure 1-Supplemental Figure 1E-G, Figure 1-Supplemental Figure 2E-G, Figure 1-Supplemental Figure 3B, D, F and insets, and Figure 2-Supplemental Figure 3). We recorded the simulation data every millisecond. Unless otherwise specified, we used a nuclear diffusion coefficient of 2 µm^2^/s, concentrations of 110 nM for Mbp1 and 150 nM for Swi6 in cells of all sizes, and 50 and 100 nM for Swi4 in small and large cells respectively, previously determined by N&B-based absolute measurements (Dorsey et al., 2018). All species were assumed dimeric to yield the total number of Mbp1, Swi4 and Swi6 dimers in small (42, 15, 57), medium (71, 37, 97) and large (131, 109, 178) simulated cells.

## Supporting information

Swi4-mEos3.2 single trajectory

Mbp1-mEos3.2 single trajectory

Swi6-mEos3.2 single trajectory

## Acknowledgements

We thank Derek McCusker for providing the mEOS3.2 plasmid. This work was supported by the National Science Foundation (PHY 1806638 to C.A.R.), the Canadian Institutes of Health Research (MOP-366608 to M.T.), the Canada Foundation for Innovation (30789 and 31072 to M.T.), and by a Canada Research Chair in Systems and Synthetic Biology (to M.T.).

---

**Figure 1-Supplemental Figure 1.**
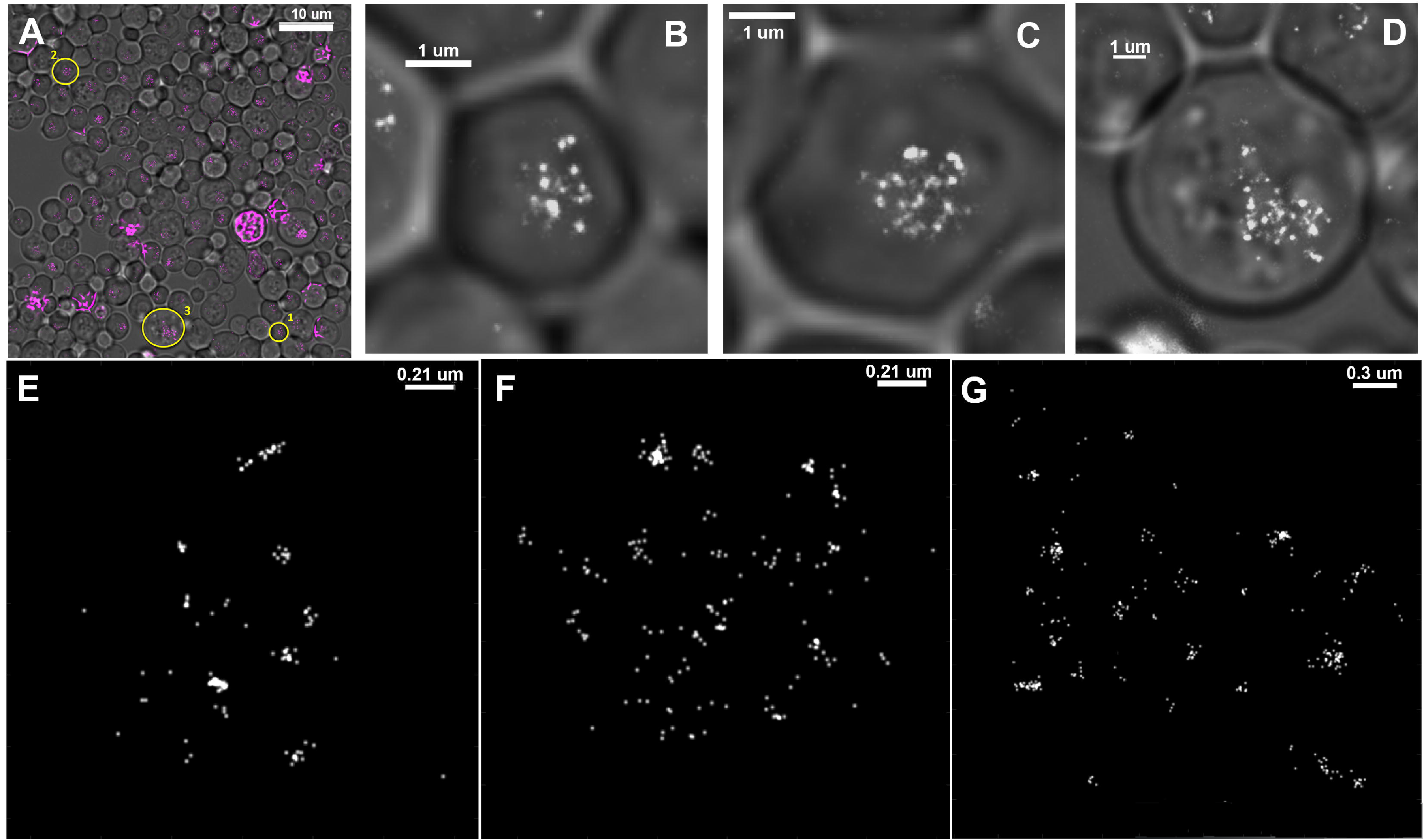
Super-resolution PALM imaging of Mbp1-mEos3.2 in fixed yeast cells. A) Composite phase contrast image of a FOV of Mbp1-mEos3.2 and the detection image (pink dots; one for each detection) output produced with Thunderstorm. The detection image is not filtered for blinking. Saturated purple intensity in the image corresponds to out of focus beads or compromised cells. Zoomed cells 1, 2 and 3 described below are indicated by red circles and numbers. Cells were grown in SC+2% glucose. Scale bar is 10 μm. B-D) Zoomed composite images of small (1), medium (2) and large (3) cells. Scale bars are 1 μm. E-F) Molecular images of the nuclei of indicated cells were created as described in the text and corrected for blinking. Scale bars are 0.21 μm in E, 0.21 μm in F and 0.30 μm in G.

**Figure 1-Supplemental Figure 2.**
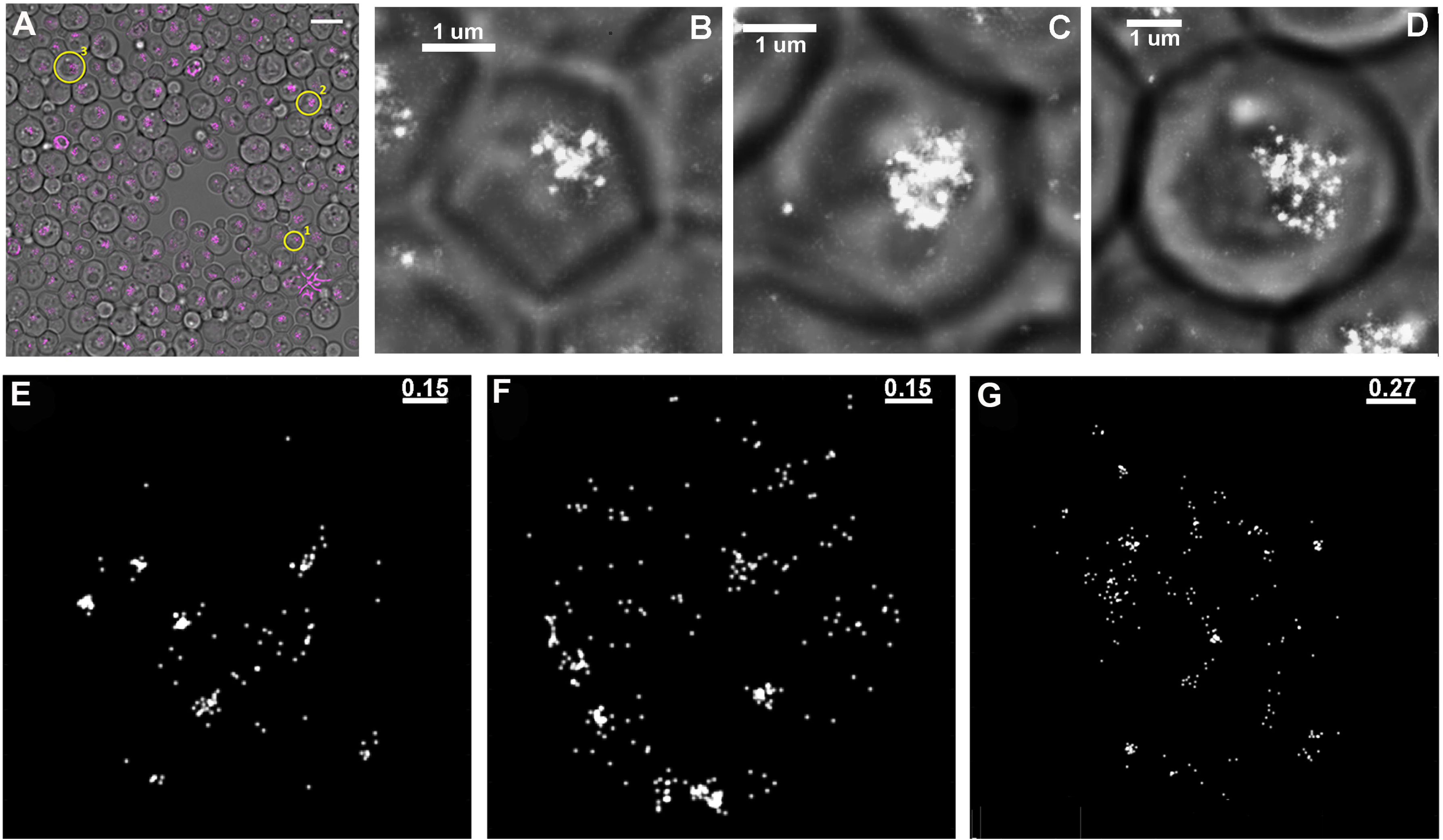
Super-resolution PALM imaging of Swi6-mEos3.2 in fixed yeast cells. A) Composite phase contrast image of a FOV of Swi6-mEos3.2 and the detection image (pink dots; one for each detection) output by Thunderstorm. The detection image is not filtered for blinking. Saturating purple intensity in the image corresponds to out of focus beads or compromised cells. Zoomed cells 1, 2 and 3 described below are indicated by yellow circles and numbers. Cells were grown in SC+2% glucose. Scale bar is 10 μm. B-D) Zoomed composite images of small (1), medium (2) and large (3) cells. Scale bars are 1 μm. E-F) Molecular images of the nuclei of cells indicated above. Molecular images were created as described in the text and corrected for blinking. Scale bars are 0.15 μm in E, 0.15 μm in F and 0.27 μm in G.

**Figure 1-Supplemental Figure 3.**
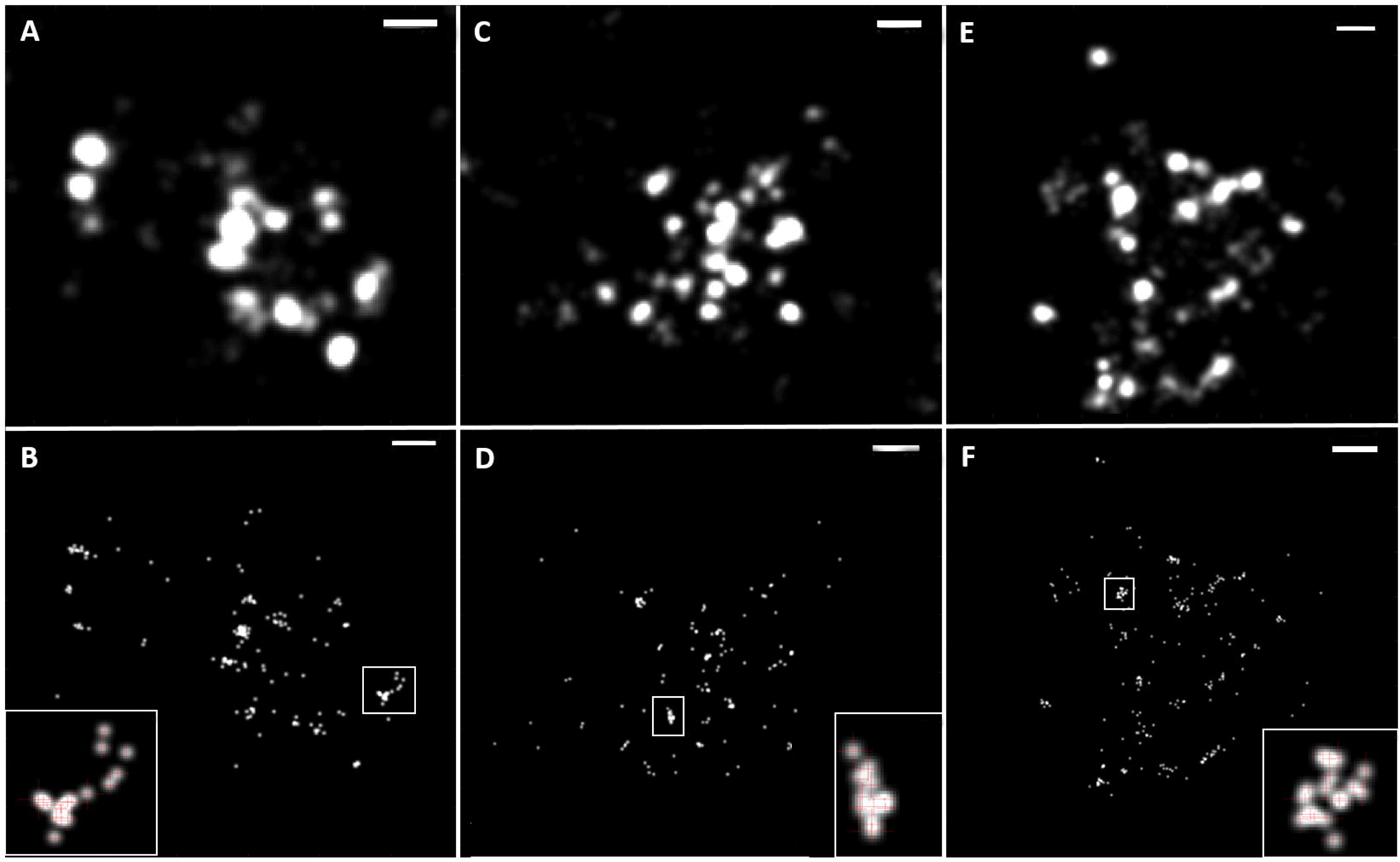
PALM images in poor nutrients. Swi4-mEos3.2 (A, B), Mbp1-mEos3.2 (C, D) and Swi6-mEos3.2 (E, F) in cells grown in SC + 2% glycerol medium. A, C, E) Detection images. Scale bars are 240 nm. B, D, F) Molecular images as described in the text and SI. Scales bars are 180, 240 and 300 nm, respectively. Shown in insets are zooms of particular clusters as indicated by the box in the larger image. Red crosses in zoomed image show the central position of each molecule ± the standard error of the mean.

**Figure 2-Supplemental Figure 1.**
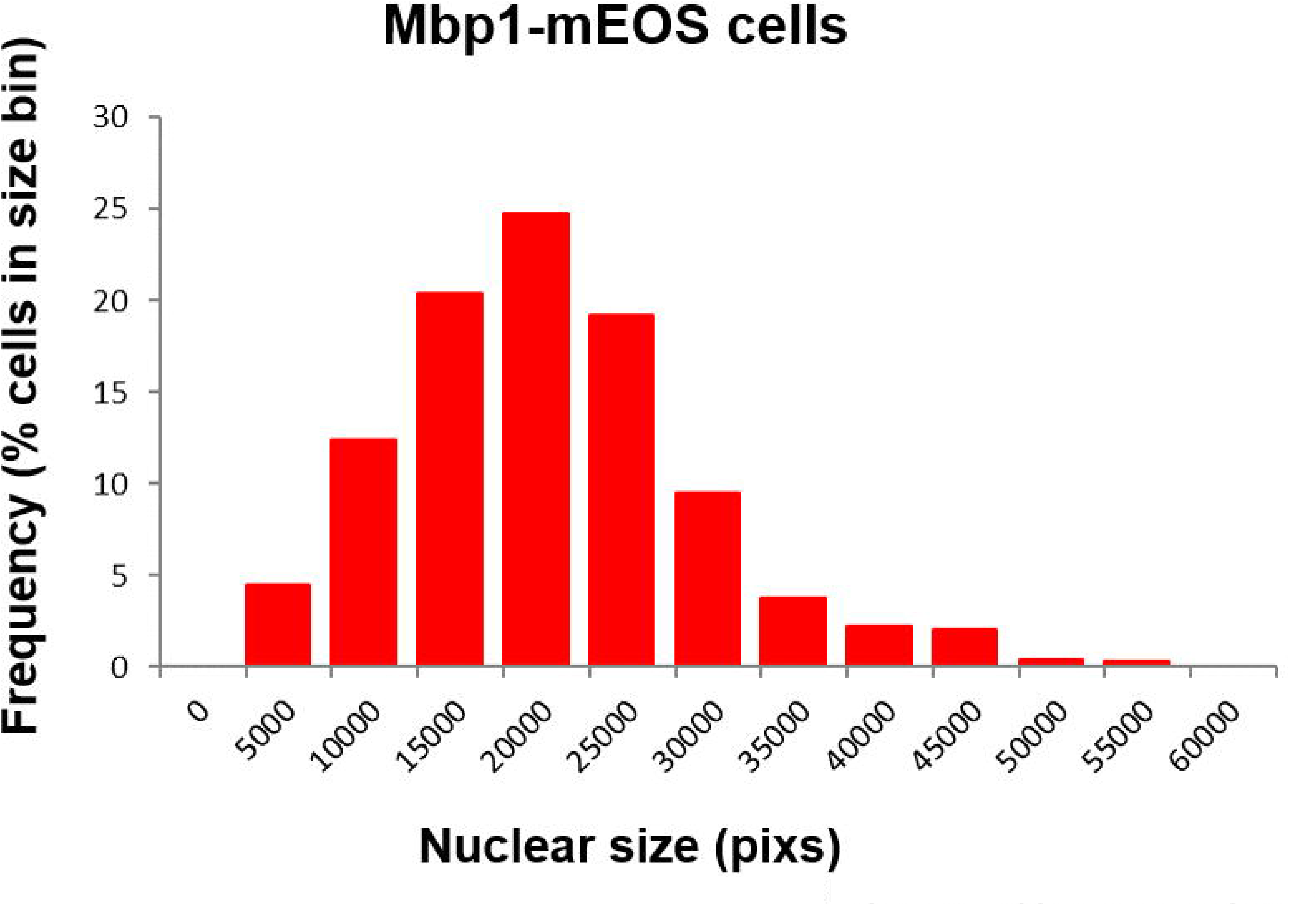
Nuclear size distribution, as determined by our masking algorithm. Distribution of nuclear sizes (in pixel units) obtained from our entire dataset of Mbp1-mEOS3.2 cells grown in SC+2%glucose medium. This distribution resembles cell size distribution of asynchronously growing cultures, therefore we used it as a proxy for cell size and in particular the size of G1/S cells (mode size, 20.000pix). Assuming spherical cells and nuclei, and the 1/7 karyoplasmic ratio (Jorgensen et al., 2007), this corresponds to a cell size of 30 fL. The Mbp1-mEOS3.2 strain was chosen in this purpose, because our analysis algorithm uses the mEOS3.2 signal to mask nuclei and, unlike Swi4, Swi6, Mbp1 nuclear signal shows no cell cycle-dependence yielding a (masked) nuclear size distribution that mirrors the entire cell size distribution.

**Figure 2-Supplemental Figure 2.**
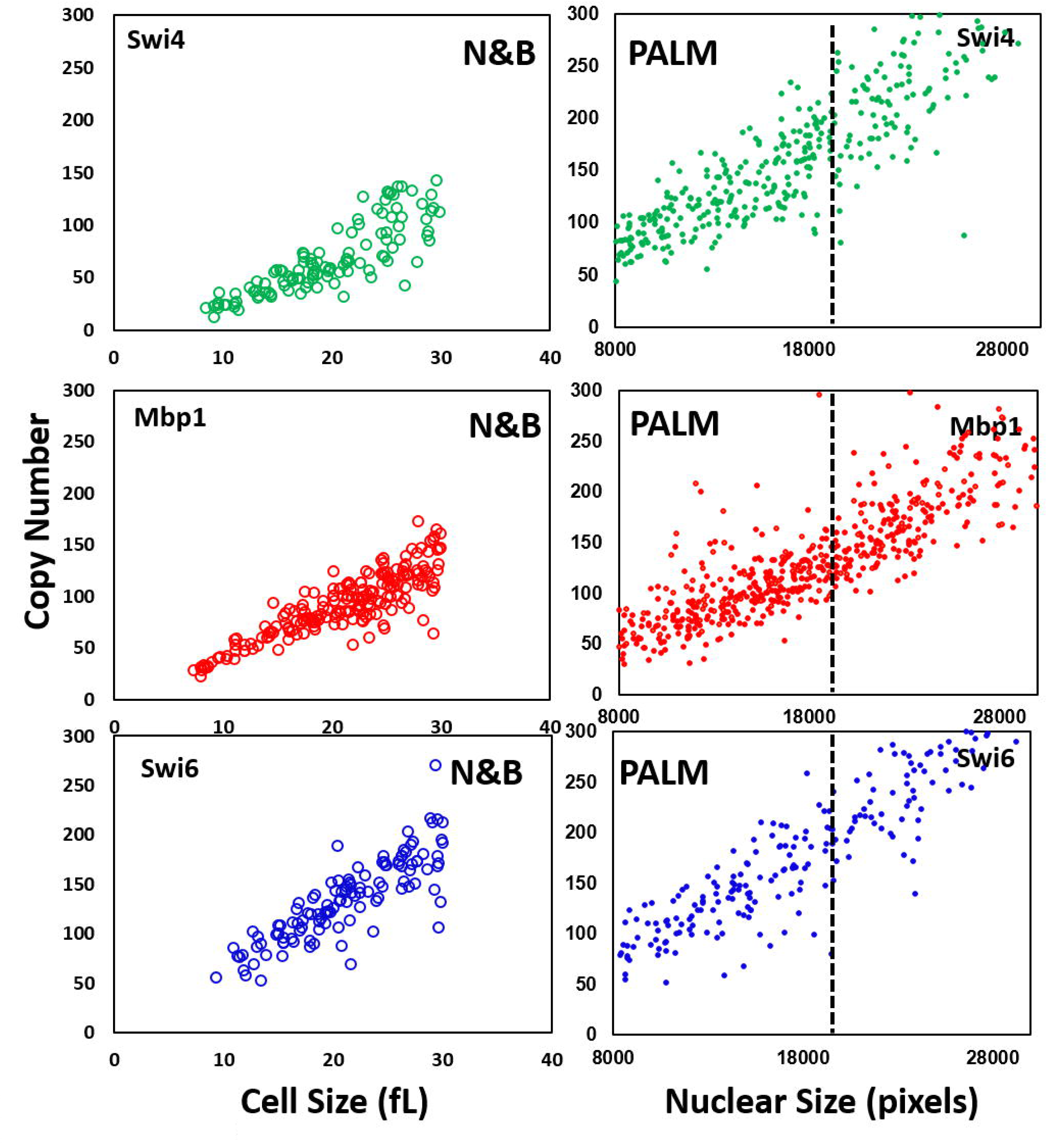
Comparison of the copy numbers of Swi4, Mbp1 and Swi6 determined by scanning number and brightness and PALM. Left panels) Copy number vs cell size (in fL) for Swi4, Mbp1 and Swi6 as indicated determined by sN&B (Dorsey et al., 2018). Only G1 cells are shown. Right panels) Copy number vs nuclear size (in pixels) for Swi4, Mbp1 and Swi6 as indicated determined by PALM. These are the data from Figure 2 A-C of the main text, in which a dotted line has been placed at the critical cell size below which the majority of cell are in G1 phase. As cells become larger, the efficiency of PALM detection decreases due to out of focus, and hence, undetected molecules. This effect is particularly clear for Mbp1 and Swi6.

**Figure 2-Supplemental Figure 3.**
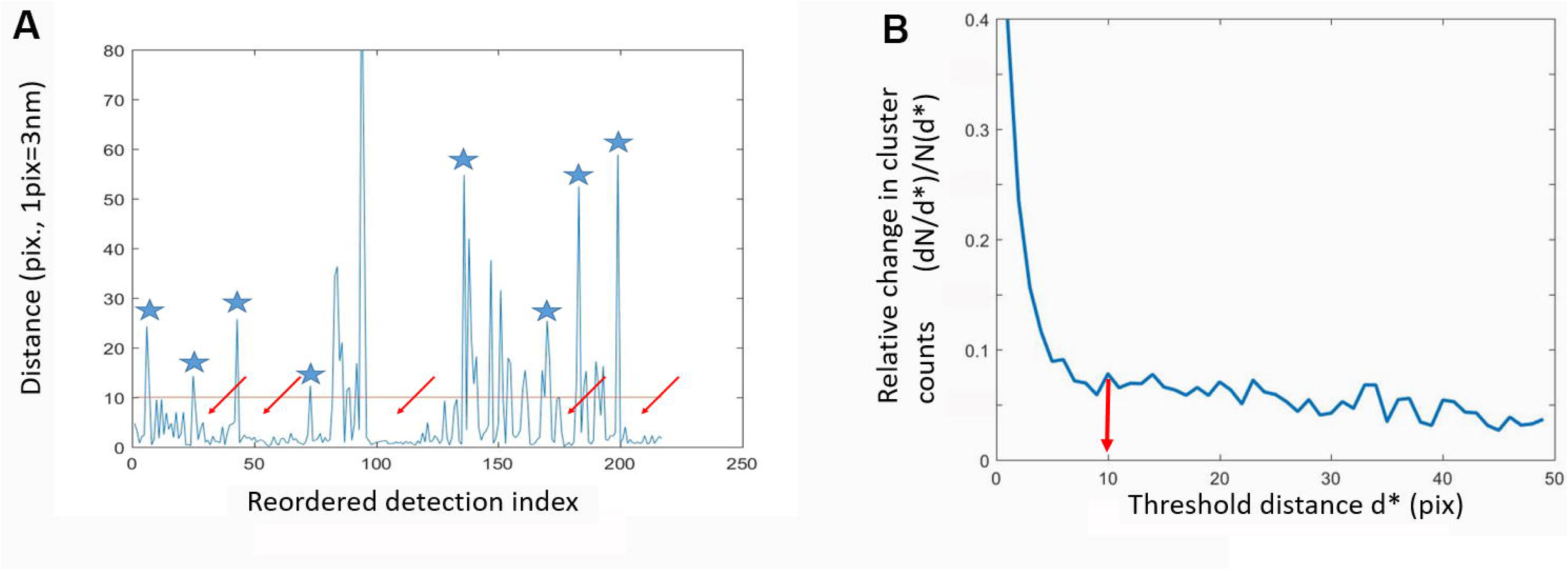
Cluster quantification algorithm. A) Distance to the next particle (vertical axis, in high resolution pixel units, 1pix=3nm) as a function of the position of the particle in the nearest-neighbor ranked list (see Methods section for details). Clusters appear as valleys (arrows), wherein inter-particle distances fall below the chosen cluster detection threshold (10pix=30nm, red line), and are separated by larger distances (blue asterisks). B) Relative variation of the number of clusters (vertical axis, cluster count change upon 3nm threshold increment over initial cluster count ratio) as a function of the specific choice of the cluster detection threshold (horizontal axis, in high resolution pixel unit, 1 pix=3 nm). Example given is for Swi4.

**Figure 2-Supplemental Figure 4.**
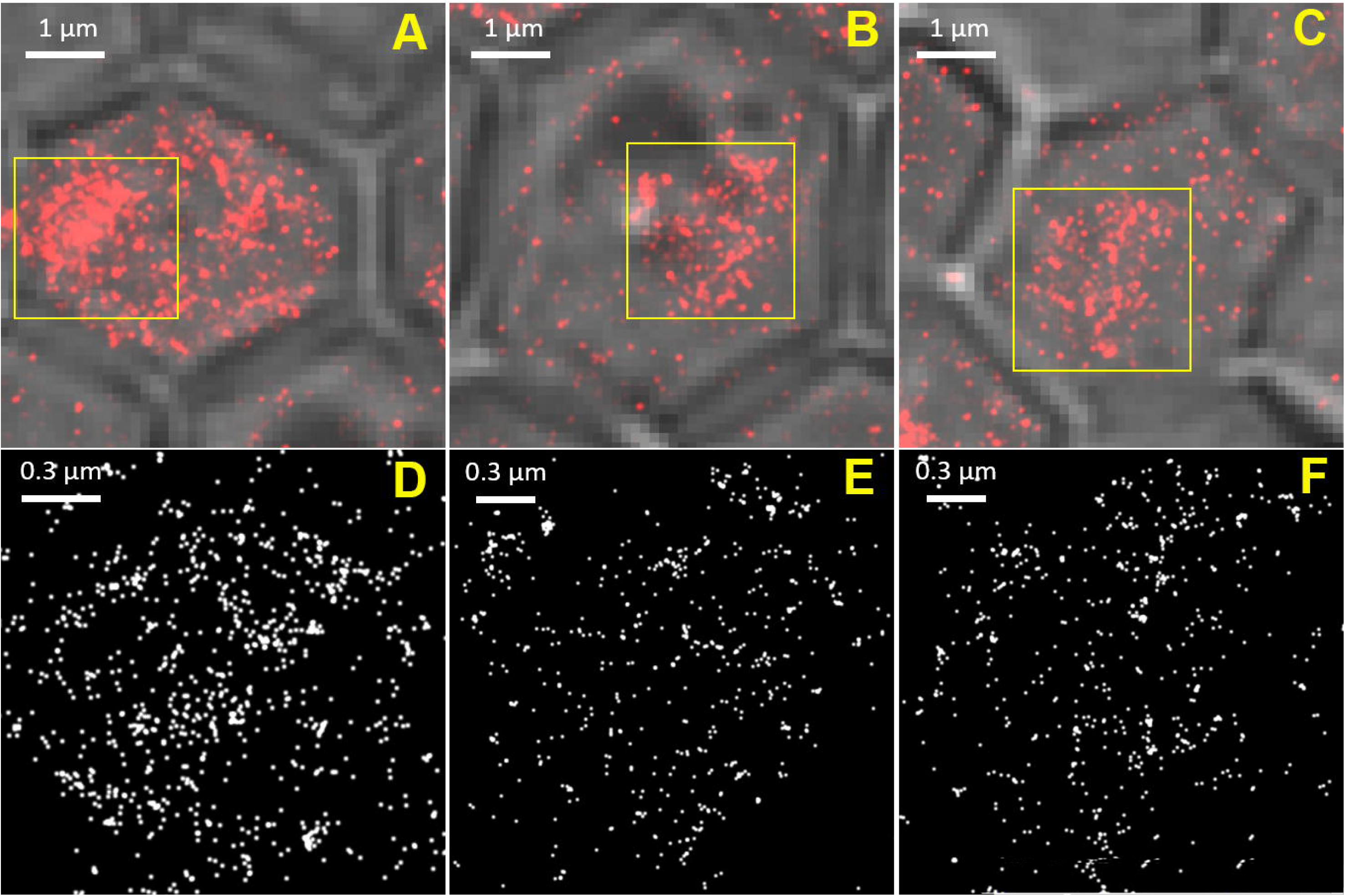
Super-resolution PALM imaging of NLS-mEos3.2-NLS in fixed yeast cells. A-C) Composite phase contrast images of three cells expressing a NLS-mEos3.2-NLS fusion protein from the GAL1 promoter. Scale bars are 1µm. D-F) Molecular images of the nuclei of the cells indicated above (from yellow squares on A-C). Each dot shows an individual NLS-mEos3.2-NLS molecule. Molecular images were created as described in the text and Methods, and corrected for blinking. Scale bars are 0.3µm.

**Figure 2-Supplemental Figure 5.**
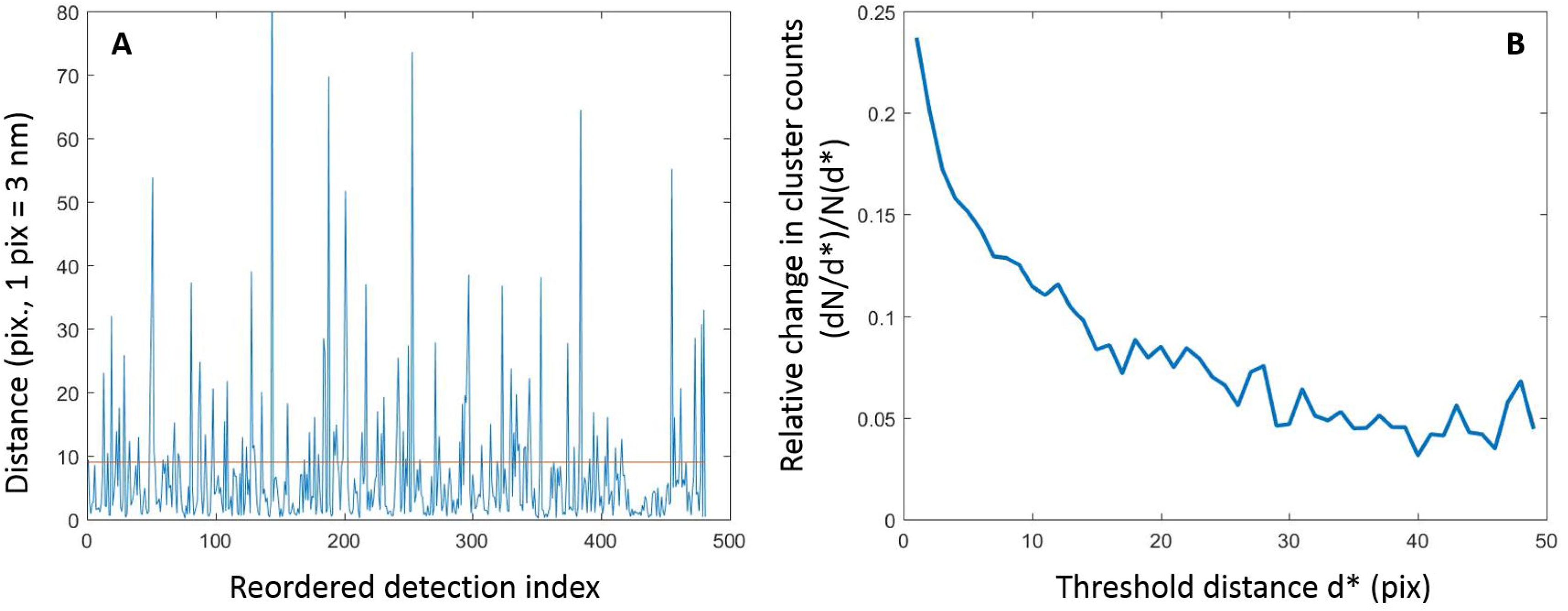
Quantification of NLS-mEOS3.2-NLS PALM data reveals no significant clustering. A) Distance to the next particle (vertical axis, in high resolution pixel units, 1pix=3nm) as a function of the position of the particle in the nearest-neighbor ranked list (see Methods section for details). The absence of large valleys in this plot (as compared to Figure 2-Supplemental Figure 3 is indicative of the absence of NLS-mEOS3.2-NLS clustering. B) Relative variation of the number of detected “clusters” (vertical axis, cluster count change upon 3nm threshold increment over initial cluster count ratio) as a function of the specific choice of the cluster detection threshold (horizontal axis, in high resolution pixel unit, 1 pix=3 nm). The absence of obvious threshold value beyond which the relative cluster counts does not change is indicative of an even distribution of NLS-mEOS3.2-NLS molecules, and of a lack of actual cluster formation.

**Figure 2-Supplemental Figure 6.**
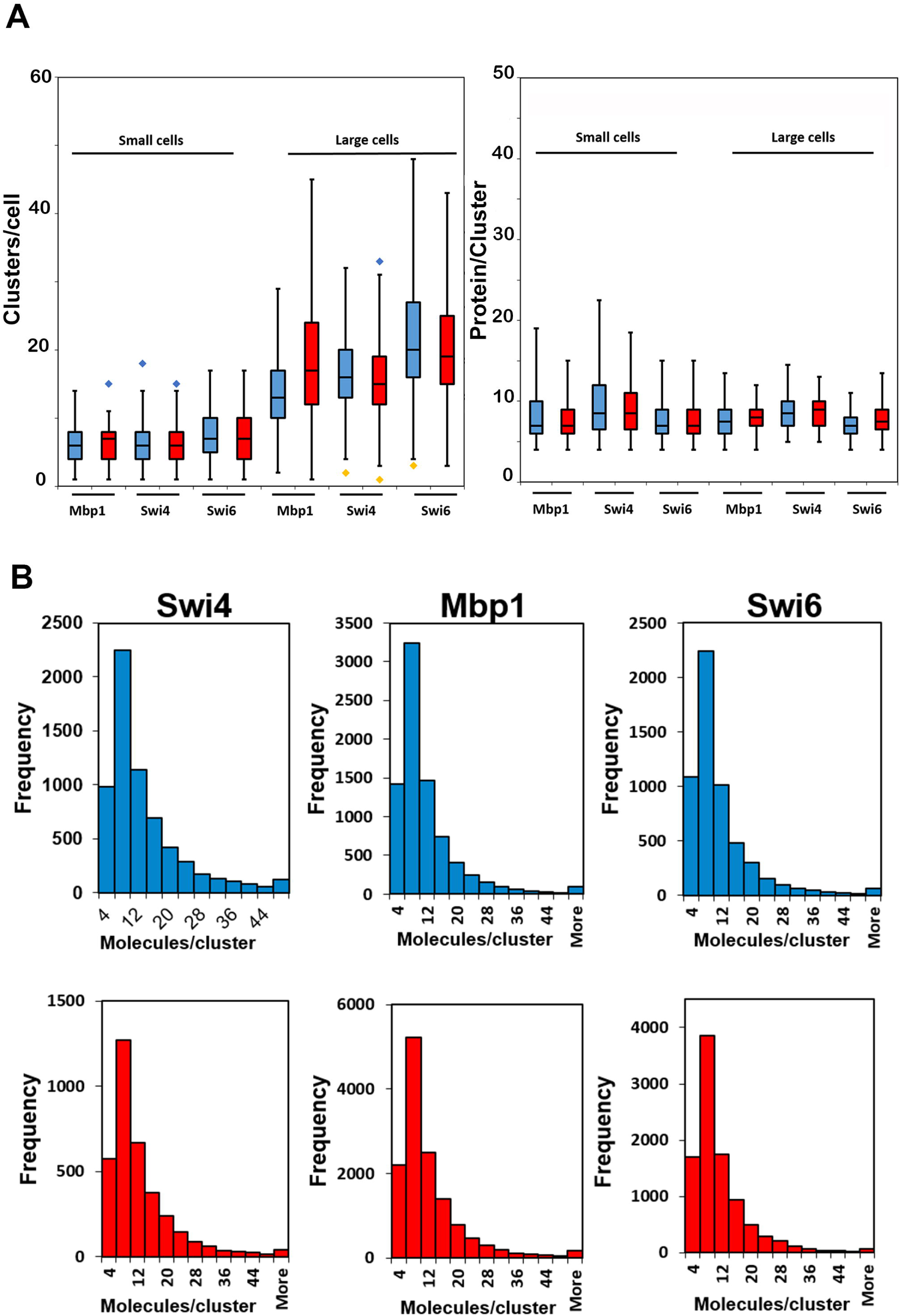
Cluster statistics are largely independent of carbon source. A) Number of clusters (left panel) and number of TF protein molecules per cluster (right panel) for Swi4, Swi6 and Mbp1 in small (<20000pix, left of each panel) and large (>20000pix, right of each panel) cells grown in glucose (blue) and glycerol (red). The Box and Whisker plots represent the distribution of values across 3-5 experiments encompassing >100 cells each for each condition, and blue and yellow diamonds represent distribution outliers. The difference in cluster counts between small and large cells was statistically significant in each condition (p-values <1e-40). There were significantly more Swi6 clusters than Swi4/Mbp1 clusters, both in large and small cells and in glucose and glycerol (p-values ranging from 10e-23 to 0.0197). B) Histograms for all clusters of Swi4, Mbp1 and Swi6 as indicated in all nuclei for cells grown in SC + 2% glucose (blue) and SC + 2% glycerol (red). The frequency unit (vertical axis) is actual cluster counts within each bin across all datasets (3-5 experiments, >100 cells each) for each condition.

**Figure 3-Supplemental Figure 1.**
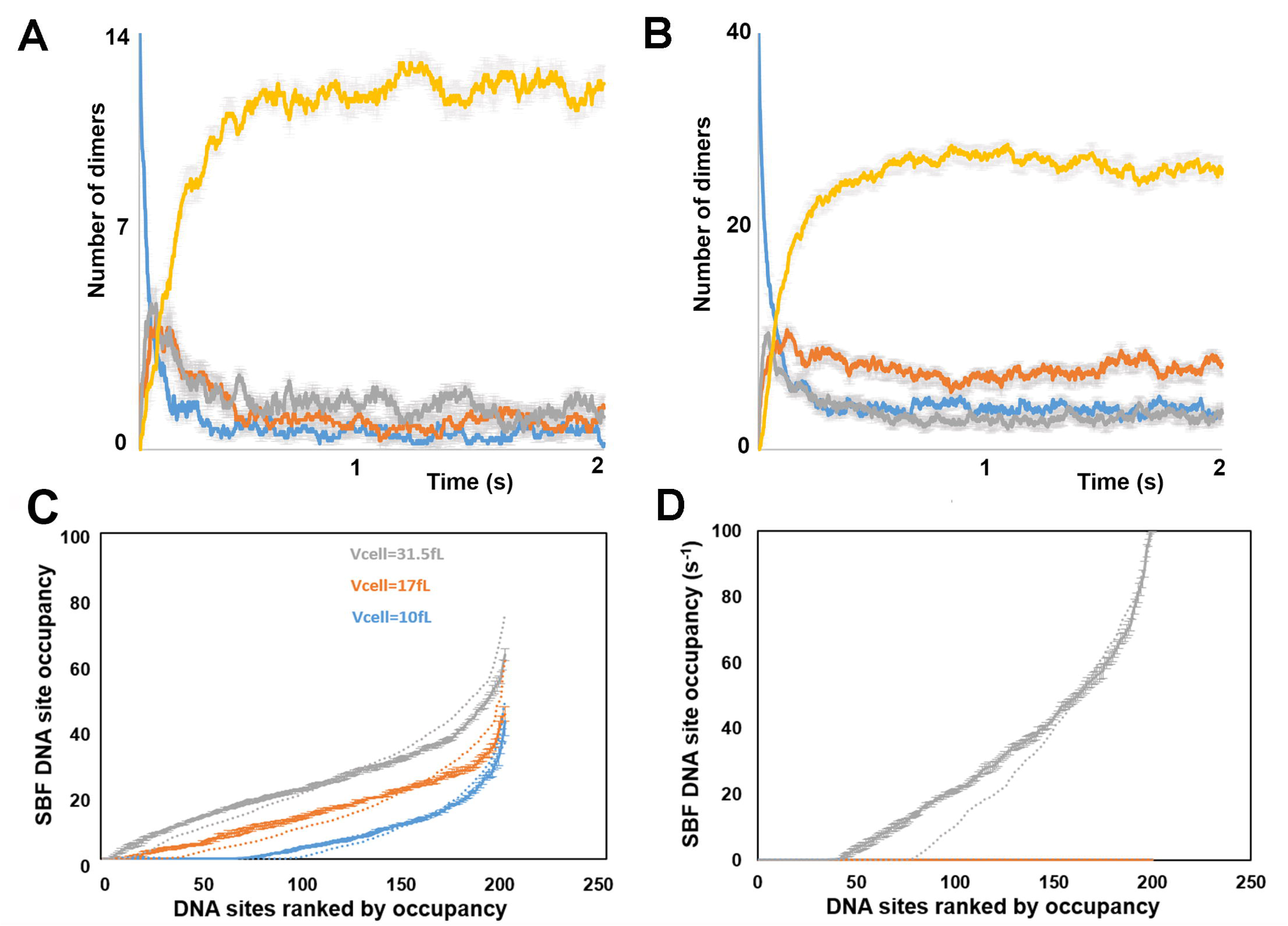
Simulated transcription factor assembly dynamics and DNA site occupancy. A and B) Typical time evolution of the number of transcription factor complexes in small (10fL) cells showing the number of each molecular species, (A) Swi4-containing species: DNA-free Swi4 (dimer) (blue), DNA-free SBF (grey), DNA-bound Swi4d (dimer) (orange), DNA-bound SBF (yellow) and (B) Mbp1-containing species: DNA-free Mbp1d (dimer) (blue), DNA-free MBF (grey), DNA-bound Mbp1d (dimer) (orange), DNA-bound MBF (yellow). Plots are for averages over 10 independent simulations. Error bars indicate the standard error on the mean at each time point for each species. C) Clustering homogenizes DNA occupancy. Promoter occupancy represented as fraction of time in the steady state (vertical axis, % of simulated time (5s) beyond 0.6s) during which each individual G1/S promoter (represented by individual dots) is occupied by SBF in small (blue), medium size (orange) and large (grey) cells. To facilitate data visualization, promoters were ranked according to increasing occupancy, which was averaged at each ranking position (rather than each individual promoter) over 5 simulations conducted in the *presence* of G1/S promoter clusters. Error bars represent the standard error on the mean. Promoter occupancy defined similarly but in the *absence* of G1/S promoter clustering is shown as dotted lines for comparison. D) Clustering improves Start synchrony. Promoter SBF-occupancy represented as fractions of time (vertical axis, %) within the last second of large time (30s) simulations in large cells (i.e., close to the G1/S transition). As in C), promoters were ranked according to increasing occupancy, which was averaged at each ranking position (rather than each individual promoter) over 5 simulations conducted in the *presence* of G1/S promoter clusters. Error bars represent the standard error on the mean. Promoter occupancy defined similarly but in the *absence* of G1/S promoter clustering (and thus, TF clustering, see Figure 3 – Supplemental Figure 1C) is shown as a dotted line for comparison. The number of promoters that are never SBF-bound during this critical time where the cell is susceptible to trigger Start at any time is 2-fold larger in the absence of clustering, indicating that clustering may improve Start synchrony.

**Figure 3-Supplemental Figure 2.**
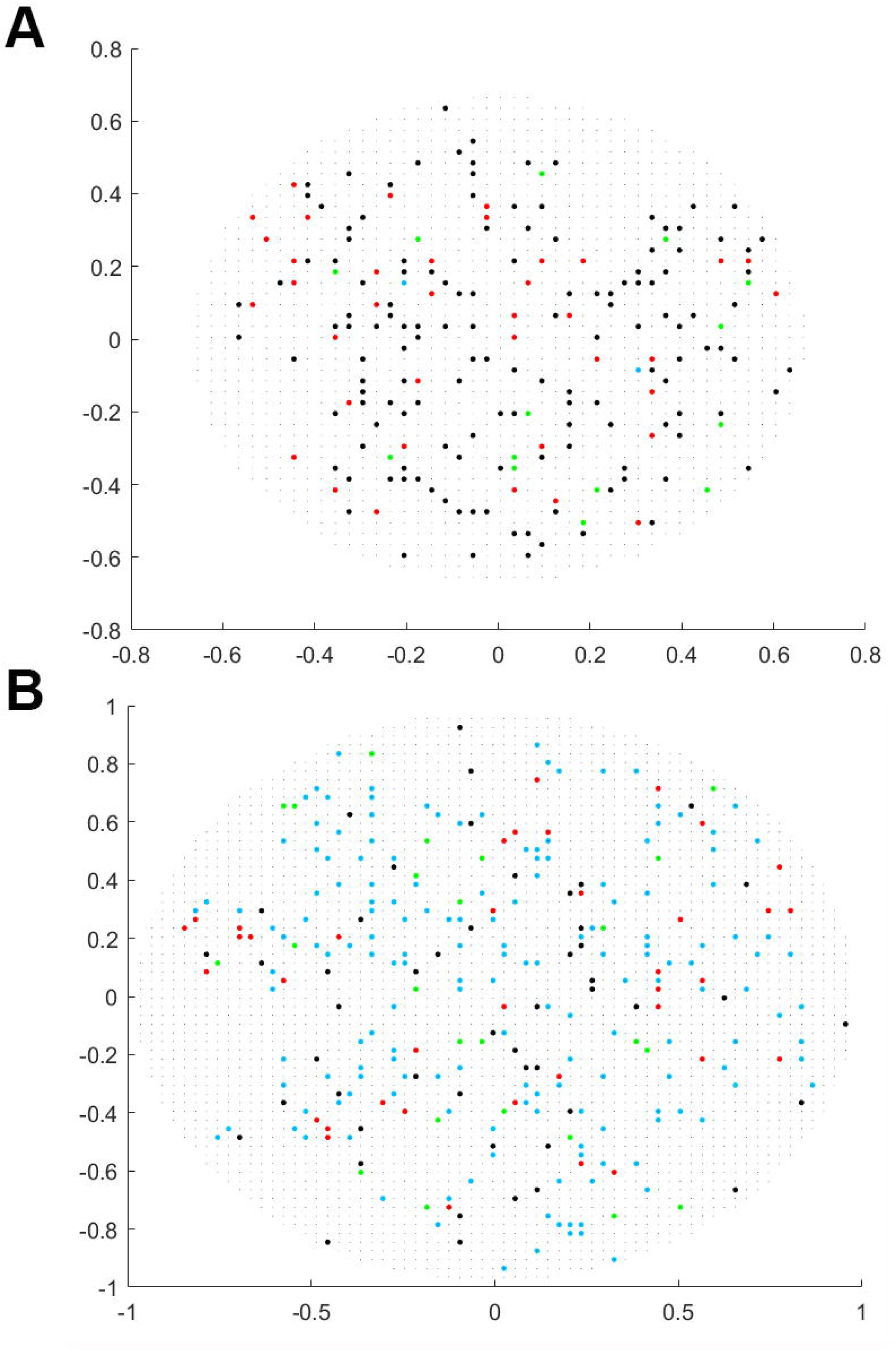
G1/S TFs do not cluster in simulations where the promoter target sites are not clustered. 2-D projections of the 3-dimensional output of a typical simulation in a small cell (A) and a large cell (B) in the absence of G1/S promoters clustering, showing Swi4 dimers (green dots), Mbp1 dimers (red), Swi6 dimers (blue) and G1/S DNA promoters (black dots) in small (10fL, left) and large (31.5fL, right) cells. Not all molecules/sites are shown since at many pixels Swi4, Mbp1, Swi6 and G1/S promoters overlap.

**Figure 3-Supplemental Figure 3.**
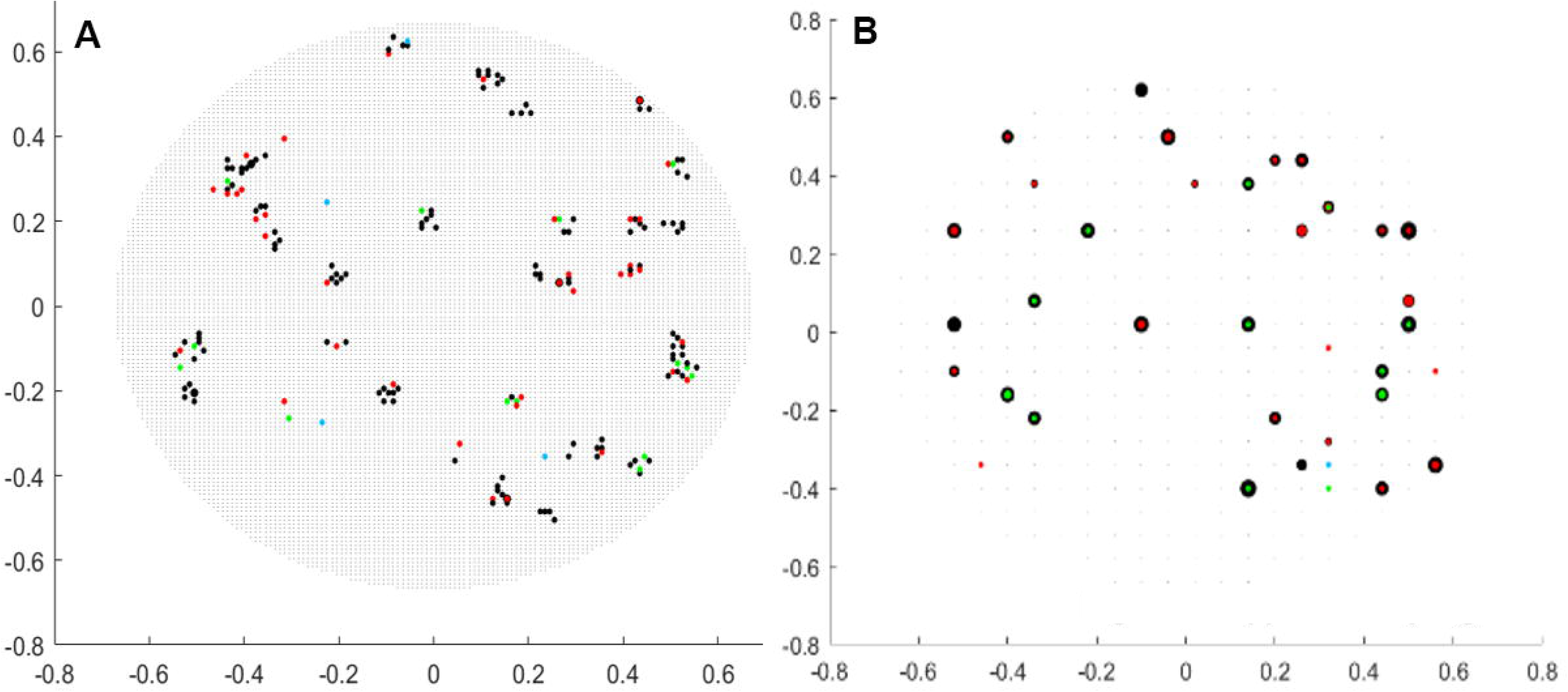
G1/S TFs clustering is independent of simulation mesh size. 2-D projections of the 3-dimensional output of a typical simulation in a small cell (10fL) using a finer (10nm, panel A) and a coarser (60nm, panel B) mesh compared to main Figure 3. As with the default 30nm mesh (Figure 3), Swi4 dimers/SBF (green dots), Mbp1 dimers/MBF (red), and free Swi6 dimers (blue) are clusters at and around G1/S DNA promoters (black dots). Dot size is proportional to the number of particles of a given type at the same mesh pixel. Not all molecules/sites are shown since at many pixels Swi4, Mbp1, Swi6 and G1/S promoters overlap, especially in the coarse simulation (B).

**Figure 3-Supplemental Figure 4.**
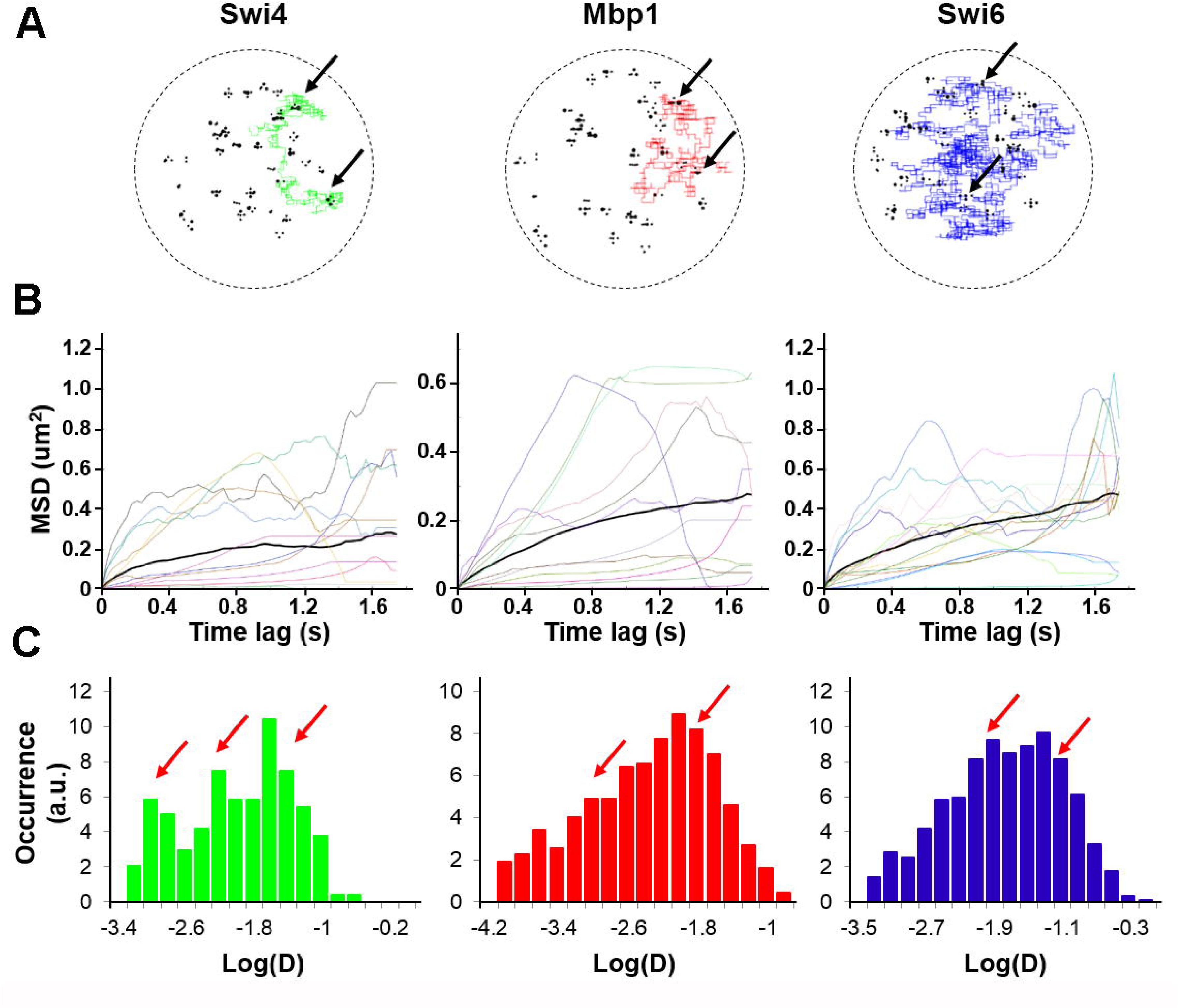
Stochastic modeling predicts a bimodal motion of individual molecules. A) Example of simulated Swi4 (dimer) (left, green), Mbp1d (middle, red), Swi6d (right, blue) single dimer trajectories in 3-dimensions, showing for each particle alternations between anomalous sub-diffusive motion confined within cluster and fast, freely-diffusive motion between clusters. Black circles represent DNA G1/S promoter target sites B) Example of MSD versus time lag curves corresponding to all individual Swi4 (left), Mbp1 (middle) and Swi6 (right) dimer trajectories (color curves) of one small cell simulation. The thick black curve represent the MSD averaged over all the trajectories for each protein. Computation of MSD curves were restricted to the steady state section of each trajectory, corresponding to times beyond 0.6s from the onset of the simulation (see Figure 3 – Supplemental Figure 1A, B). C) Histograms of single trajectory diffusion coefficients extracted from the slope of linear fits of the first 4 points of individual MSD curves from panel B. Data was gathered from 5-7 independent simulations. Red arrows indicate the approximate positions of main peaks underlying the distribution. We note that simulations predicted a significant fraction of quasi-immobile molecules with effective diffusion coefficient lower than 0.001. We did not observe this fraction in experiments, possibly due to instrument jitter which sets a lower bound to the slowness of motions we can quantify.

**Figure 4-Supplemental Figure 1.**
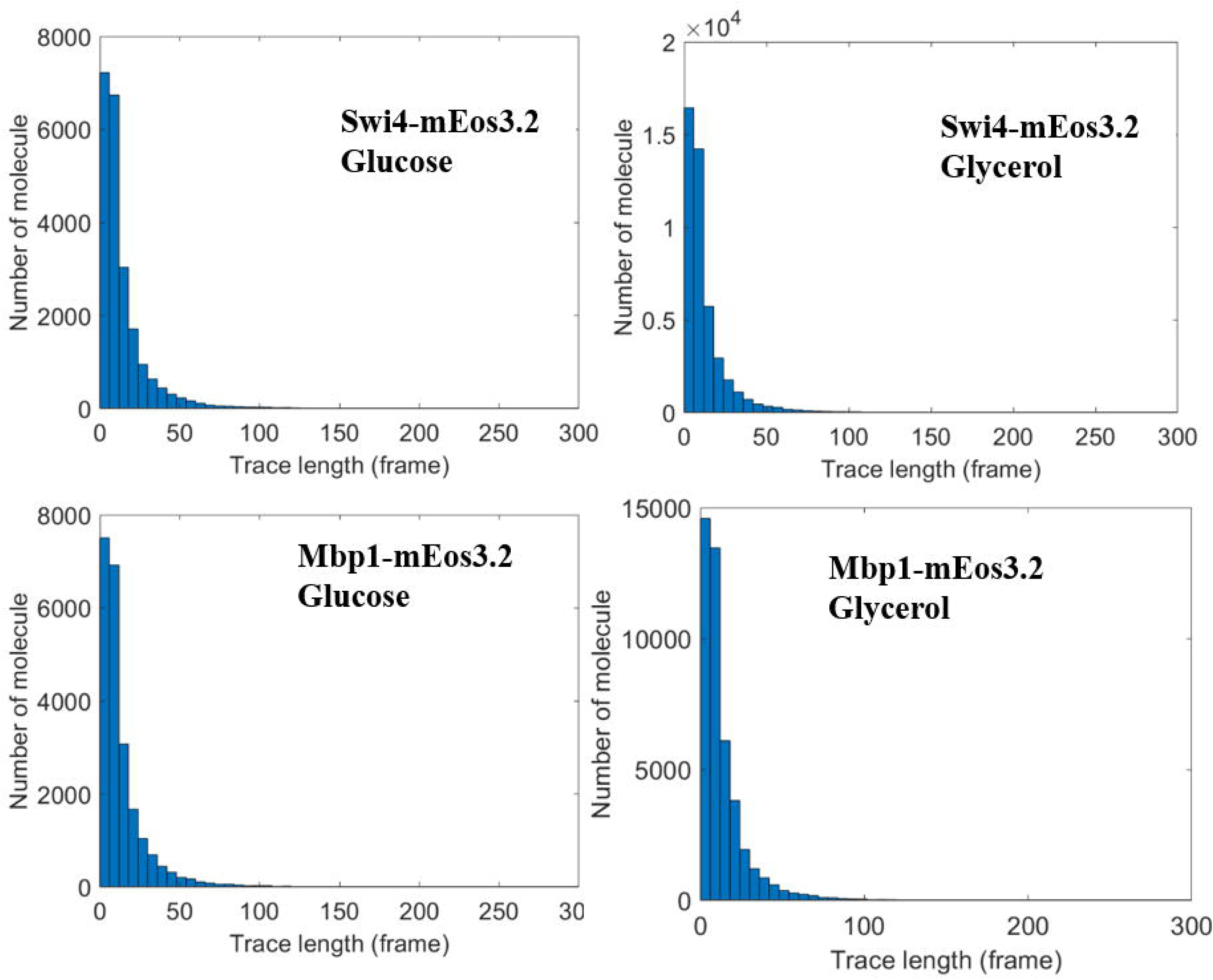
Trajectory length distributions for sptPALM on mEos3.2-Swi4 and Mbp1 in live cells. A, B) mEos3.2-Swi4 in cells grown of SC+2% glucose (A) and SC+2% glycerol (B). C, D) mEos3.2-Mbp1 in cells grown of SC+2% glucose (C) and SC+2% glycerol (D).

**Figure 4-Supplemental Figure 2.**
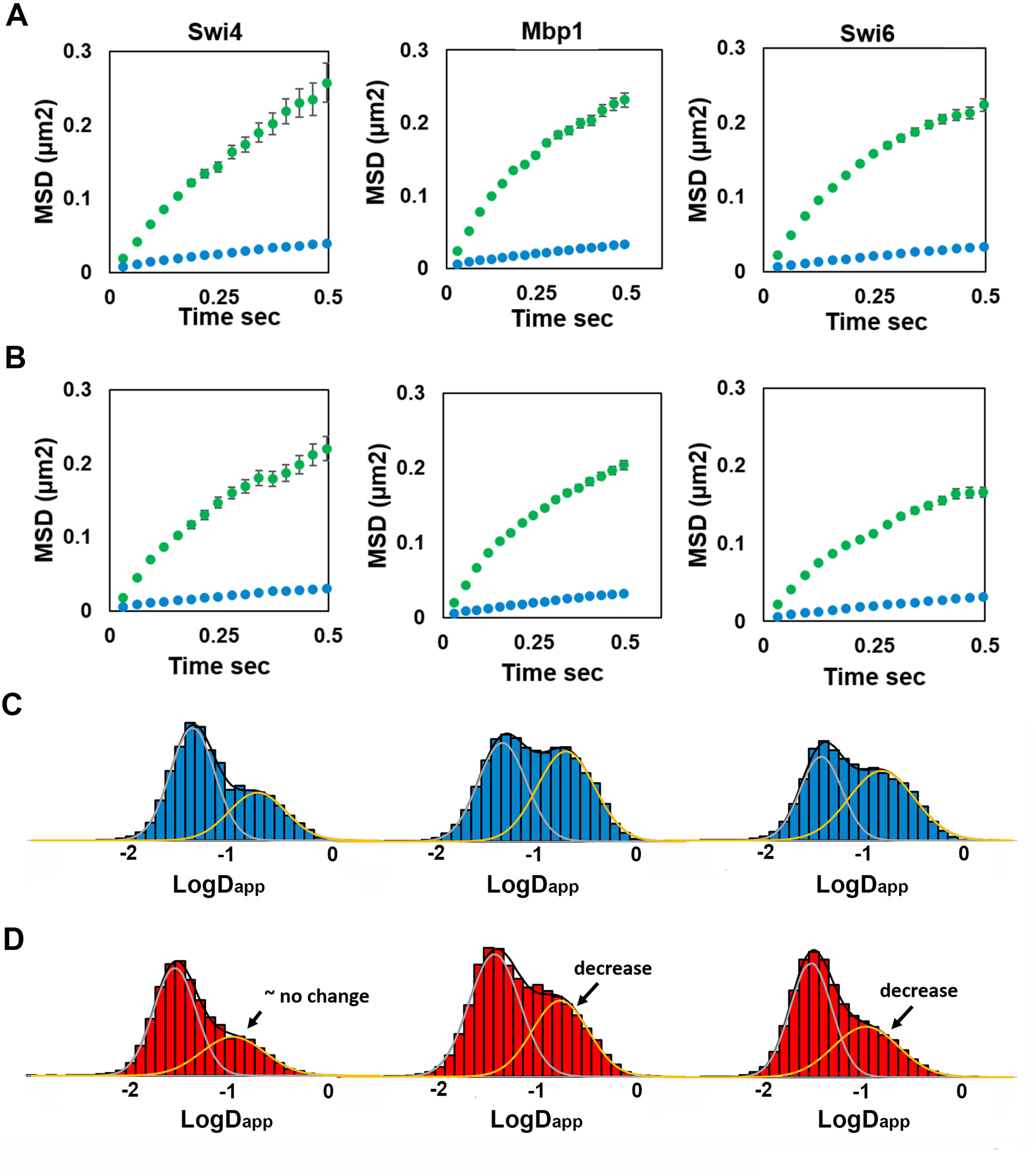
Two regimes of TF motion are evident in rich and poor carbon sources. A and B) Plots of the average mean squared displacements for all trajectories of mEos3.2-tagged Swi4, Mbp1 and Swi6 are shown for SC+2% glucose (A) or SC+2% glycerol (B) medium. Two classes of motion are apparent in all cases, one fast (green) and one slow (blue). C and D) Distributions of diffusion coefficients from two component analysis of complete individual molecule trajectories for mEos3.2-labeled Swi4, Mbp1 and Swi6 (left, middle and right) in SC+2%glucose (C) and SC+2%glycerol (D) medium. A decrease in the fraction of the fast component is apparent for Mbp1 and Swi6 in the poor carbon source.

**Figure 4-Supplemental Figure 3.**
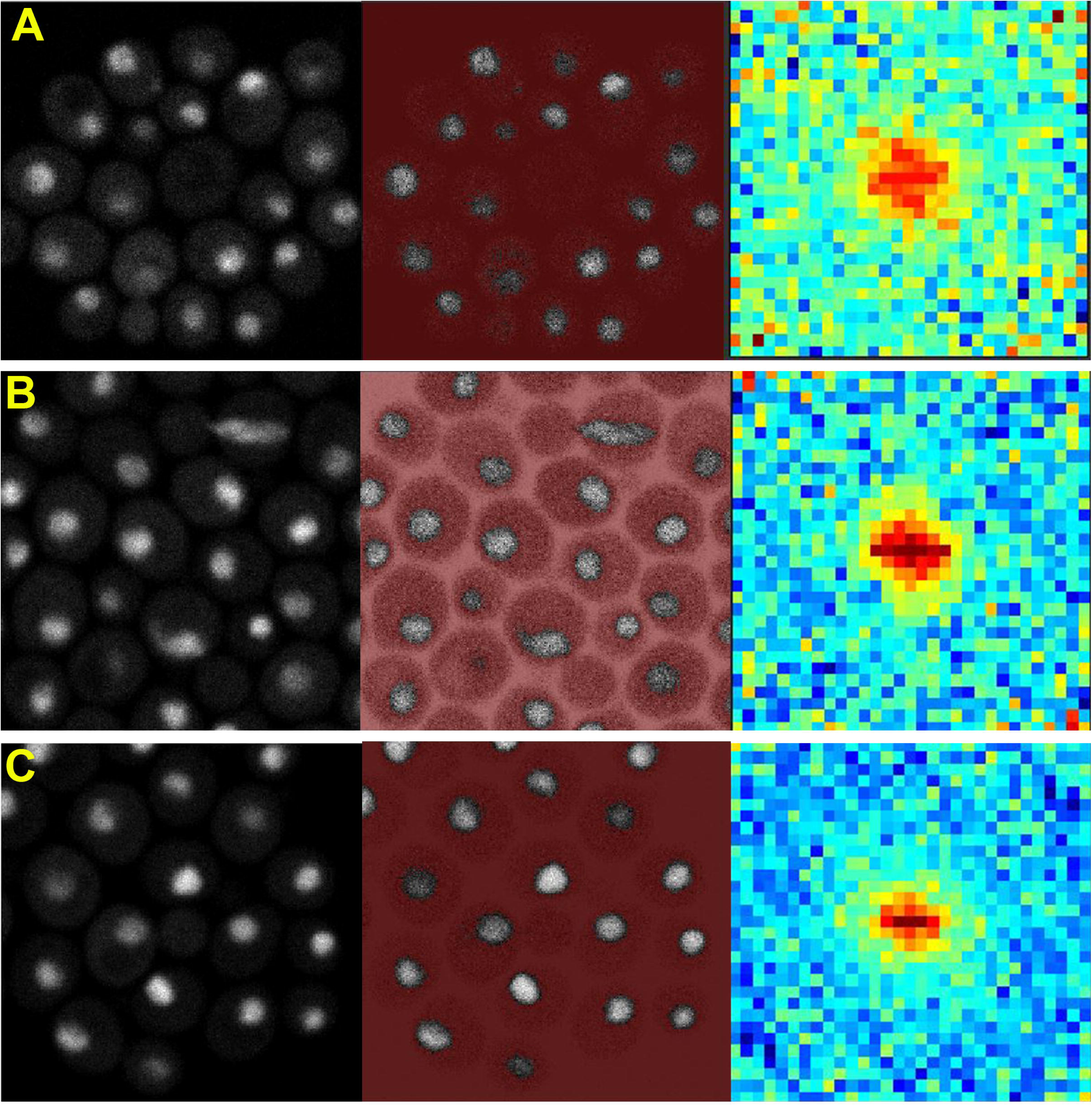
ARICS Analysis of Diffusion of G1/S TF-GFP fusion constructs. Strains expressing GFPmut3 fusion constructs of Swi4, Mbp1 and Swi6 from their natural loci were as previously described (Dorsey et al., 2018). ARICS analysis (Hendrix et al., 2016) allows RICS analysis of arbitrary regions of an FOV. ROI were selected based on intensity threshold values that selected only pixels in the nuclei. Spatio-temporal correlation of the intensity values was carried out as described previously (Digman et al., 2013). Left panels correspond to grey scale intensity images. Middle panels correspond to arbitrary thresholding in which only grey pixels are analyzed and red pixels are eliminated. Right panels show the RICS image. A) Swi4-GFPmut3, B) Mbp1-GFPmut3 and C) Swi6-GFPmut3. Cells were grown in SC+2% glucose medium. Note that RICS analysis is limited to timescales faster than ∼50 ms, with any slower dynamics appearing to be immobile on the RICS timescale. Thus, these spatio-temporal correlation patterns are a signature of TF motion at shorter timescales.

**Figure 4-Supplemental Figure 4.**
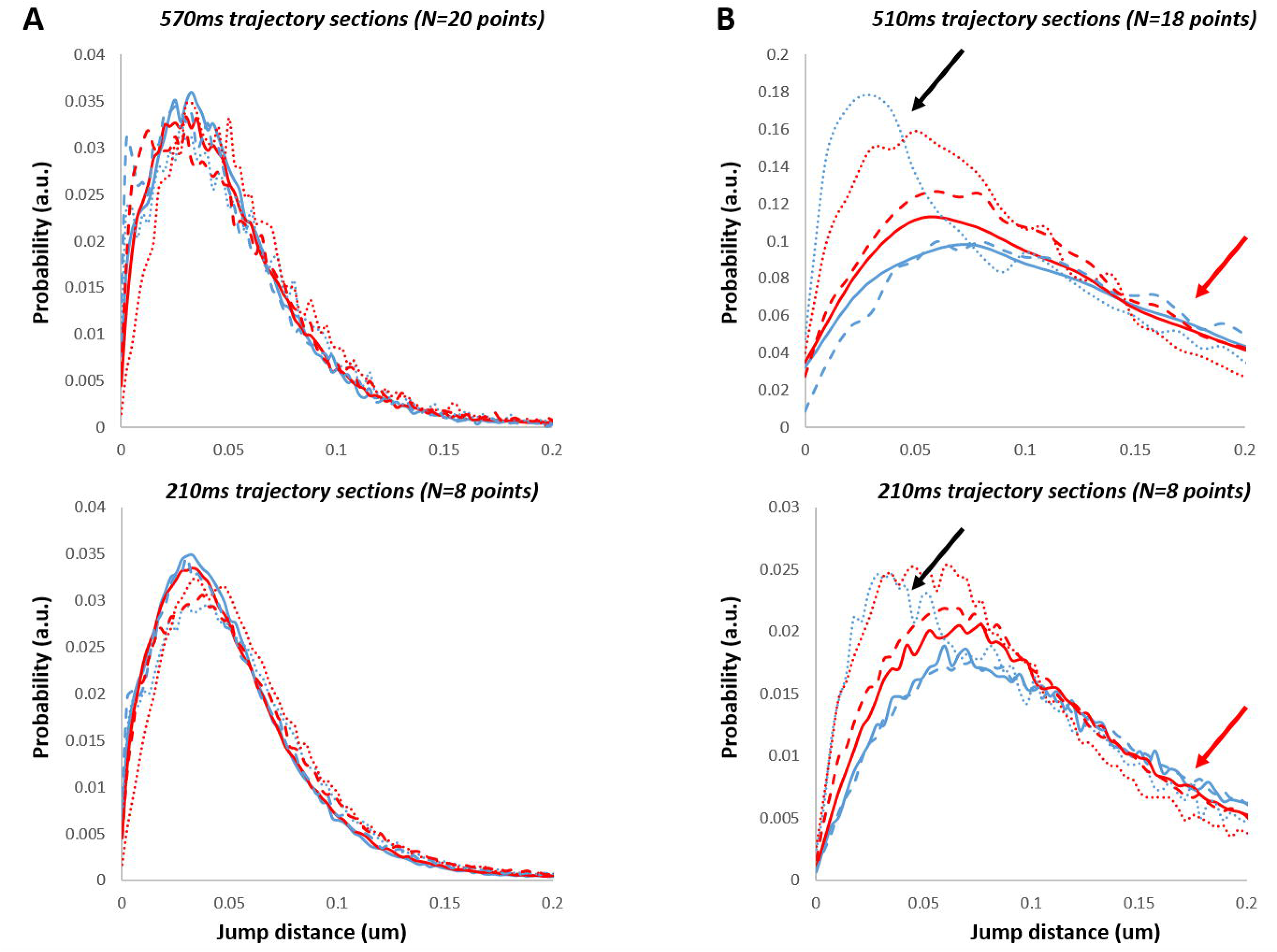
Jump size distribution analysis of spt-PALM experiments. JDD analysis is shown for Swi4 (dotted lines), Mbp1 (filled lines) and Swi6 (dashed lines) in (A) fixed cells and (B) live cells grown in SC+2%glucose medium (blue) and SC+2% glycerol medium (red). Top panels show distributions of jump distances over large times (570ms and 510ms respectively for fixed and live cells) while bottom panels show distributions of jump distances over shorter times (210ms, 8-points trajectories for both fixed and live cells). Red arrows (in B) point to the regions of the live cells distributions which show less steep drop in probability with jump distance than the trajectories from fixed cells (A). This deviation indicates long jumps that exceed confined diffusion (marked by black arrows).

**Video 1. Example trajectory of mEos3.2-Swi4 in a live yeast cell nucleus.**

**Video 2. Example trajectory of mEos3.2-Mbp1 in a live yeast cell nucleus.**

**Video 3. Example trajectory of mEos3.2-Swi6 in a live yeast cell nucleus.**

